# The Zelda Interactome Reveals Diverse Co-factors Essential for Pioneer Factor-Mediated Zygotic Genome Activation

**DOI:** 10.64898/2026.02.23.707495

**Authors:** Xiao-Yong Li, Michael B. Eisen

## Abstract

Zelda (Zld) is the master pioneer transcription factor essential for zygotic genome activation (ZGA) during the *Drosophila* maternal-to-zygotic transition (MZT). While Zld is known to promote chromatin accessibility at early enhancers and possess multiple conserved functional domains, the specific protein complexes it recruits to execute its functions remain poorly defined. Using an optimized immunoprecipitation and mass spectrometry (IP-MS) pipeline in Stage 4–5 embryos, we identified a diverse Zld interactome. This repertoire includes the coactivators dCBP and Fsh (the *Drosophila* Brd4 ortholog), subunits of major nucleosome remodeling complexes(e.g. PBAP/BAP, NURF, FACT), RNA polymerase II (RNAPII), the corepressor Smrter (Smr), and the Tousled-like kinase (Tlk). We confirmed the enrichment of ten key factors at Zld-bound regions using CUT&RUN. Notably, our results demonstrate that RNAPII associates not only with Zld-bound promoters but also with distal Zld-binding sites. This finding, which contrasts with previous ChIP-based studies, suggests that Zld facilitates transcription by actively scaffolding or pre-recruiting the transcriptional machinery at enhancers. Functional analyses revealed that dCBP and Fsh are required for Zld-mediated activation, while RNAi knockdown of *smr* led to the broad derepression of Zld-target genes, suggesting that Smr modulates Zld activity to prevent premature expression. Finally, we show that Tlk, a kinase typically associated with DNA replication, directly interacts with a specific Zld domain *in vitro* and co-localizes with Zld *in vivo*. This suggests a novel mechanism by which Zld may coordinate transcriptional activation with the rapid mitotic cycles of early embryogenesis. Collectively, our findings provide a comprehensive map of the Zld interactome and reveal how a pioneer factor integrates diverse chromatin and transcriptional regulators to orchestrate zygotic genome activation.

## INTRODUCTION

In multicellular organisms, precise spatial and temporal gene expression is governed by enhancers, regulatory DNA sequences harboring binding sites for sequence-specific transcription factors(TFs) [1–3]. Most enhancers utilize unique combinations of these sites to integrate diverse developmental cues into specific transcriptional outputs. Within the nuclear chromatin context, distinct TFs may serve specialized roles in enhancer activity.

Eukaryotic genomes are organized into chromatin through the assembly of nucleosomes [4], and inactive enhancers, in particular, are characterized by high nucleosome occupancy *in vivo* [5–9]. This wrapping of DNA around histone octamers in nucleosomes can occlude DNA binding motifs and restrict TF binding through steric hindrance. Consequently, many TFs are unable to access their motifs embedded in nucleosomes *in vitro* [10,11], and bind predominantly to accessible sites *in vivo* [12–15]. However, a specialized class of TFs termed “pioneer factors” possess the intrinsic ability to bind motifs embedded in nucleosomes *in vitro* and within closed chromatin *in vivo* [16,17]. Upon binding, such factors are able to promote local chromatin accessibility, facilitating the binding by other factors to achieve transcription activation. Pioneer factors are thus essential for establishing the initial regulatory landscape during zygotic genome activation (ZGA) and for reshaping gene networks during developmental processes such as cell fate determination and lineage specification, as well as cell reprogramming [16–22].

Zld is a master pioneer factor essential for zygotic genome activation in *D. melanogaster* [23]. It binds broadly across the genome, targeting the majority of zygotic enhancers [24,25]. Notably, Zld binding occurs very early - often before the onset of robust transcription and prior to the recruitment of non-pioneer patterning factors such as Dorsal (Dl) and Caudal (Cad) [24,26]. Zld is required for histone depletion, increased chromatin accessibility, and histone acetylation at zygotic enhancers [7,8,27–31]; and it exerts a significantly more profound effect on chromatin accessibility at enhancers than typical embryo patterning factors [8,31]. By establishing accessible chromatin, Zld facilitates the subsequent binding of other factors and the activation of zygotic genes [7,8,29,31–34]. Additionally, recent studies suggest Zld may also function by forming nuclear hubs that increase the local concentration of other transcriptional regulators [34–37].

Beyond sequence-specific TFs, enhancer function requires a diverse repertoire of factors, including coactivators, chromatin modifiers, and nucleosome remodelers [38–48]. To date, however, the specific protein complexes that collaborate with pioneer factors, such as Zld, to execute their functions remain poorly defined. Given Zld’s role in actively promoting chromatin accessibility, we anticipated that it interacts with specific histone modifiers and nucleosome remodelers, as has been observed for other pioneer factors [20,22]. Structurally, Zld contains multiple conserved domains, including a putative activation domain [49] and a domain with repressive activity [50]. These domains likely recruit the basal transcription machinery, coactivators, and potentially unknown regulators to coordinate transcription during the rapid, synchronous nuclear divisions of early development.

In this study, we utilize an optimized immunoprecipitation and high-resolution mass spectrometry (IP-MS) approach to provide a comprehensive proteomic and genomic map of the Zld interactome during the *D. melanogaster* maternal-to-zygotic transition (MZT). Our findings reveal that Zld recruits a diverse repertoire of co-factors, including coactivators, nucleosome remodelers, RNA polymerase II and its associated factors, the corepressor Smarter (Smr), and the Tousled-like kinase (Tlk). We performed CUT&RUN analysis for ten of these factors, demonstrating their association with Zld binding sites in embryos. Furthermore, we investigated the functional relationships between Zld and several enriched factors. We found that the coactivators dCBP (encoded by *nej*) [51] and Fsh (encoded by *fs(1)h*) [52,53]—the latter being the ortholog of the mammalian BET (bromodomain and extra-terminal domain) factor Brd4 [54]—are essential for the transcription of Zld target genes. We also found that Smr [55,56] exerts a broad repressive effect on Zld target gene transcription. Finally, based on *in vitro* binding assays, we found that Tlk, a cell-cycle regulated kinase with activity peaking during S-phase [57], interacts directly with a specific Zld domain. This suggests a novel mechanism by which Zld may coordinate transcriptional activation with the rapid mitotic cycles during early embryogenesis. Collectively, these results elucidate how Zld integrates chromatin remodeling with the active recruitment of the transcriptional apparatus to orchestrate the initial wave of zygotic genome activation.

## RESULTS

### Identification of the Zld interactome via optimized IP-MS

To identify protein factors associated with Zld in embryos, we developed an optimized IP-MS approach based on established protocols [38–41,58,59](Fig. 1A–B). To ensure efficient fixation, crosslinking was performed by homogenizing embryos in a formaldehyde-containing buffer followed by an extended incubation. For lysate preparation from isolated nuclei, we employed only mild sonication to avoid the disruption of protein complexes. Full solubilization was achieved via Benzonase treatment; by completely digesting DNA and RNA, this step prioritized the identification of factors associating with Zld through protein-protein interactions rather than tethering via nucleic acids. Following immunoprecipitation, the isolated proteins were digested with trypsin, and the resulting peptides were analyzed by high-resolution LC-MS/MS. Data analysis was performed using MaxQuant with label-free quantification (LFQ) [60], followed by statistical analysis in Perseus [61].

**Fig. 1.**
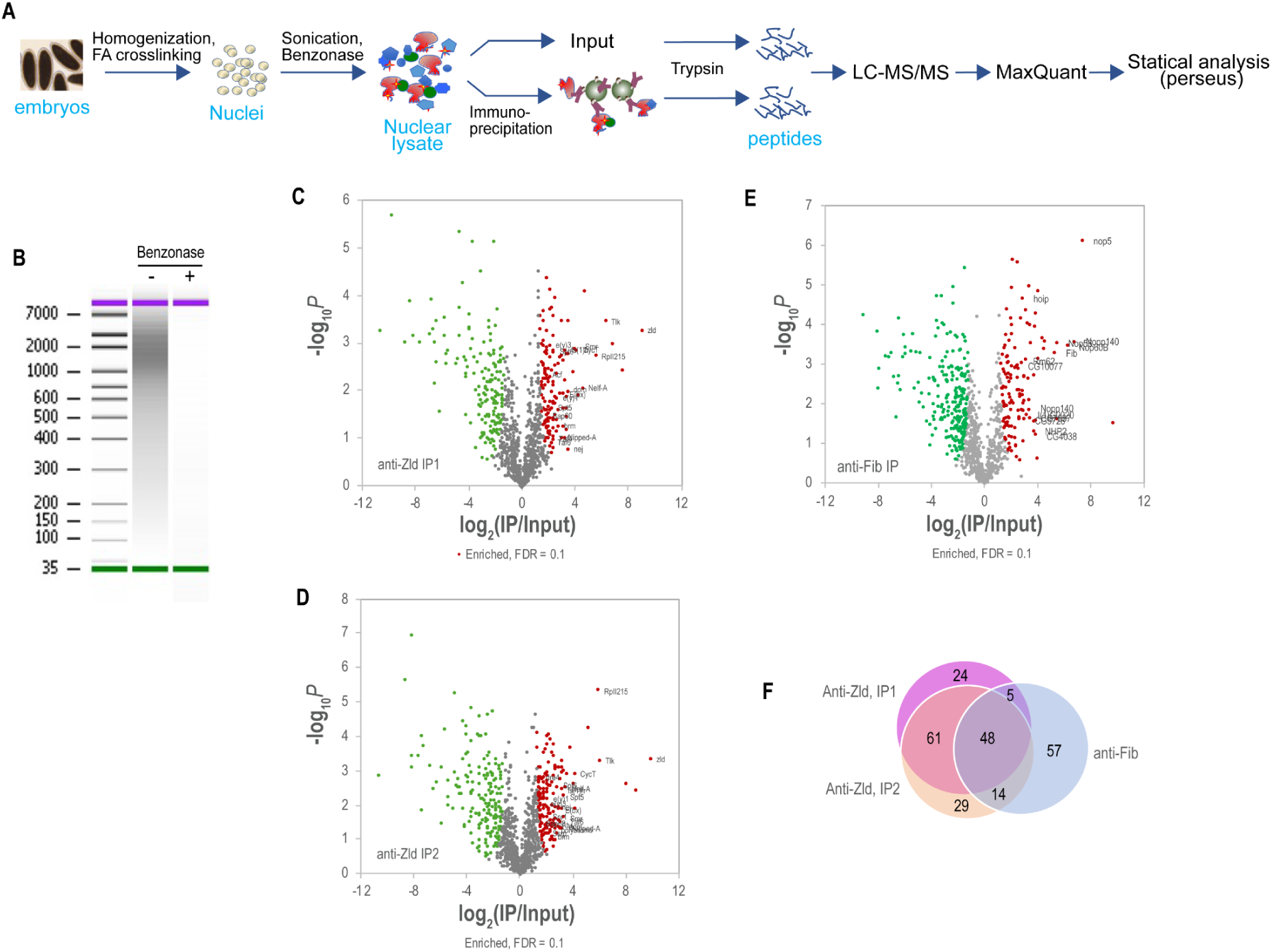
Identification of the Zelda (Zld) interactome in early D. melanogaster embryos. A) Schematic of the IP-MS workflow. Embryos were crosslinked, nuclei isolated, and chromatin solubilized with Benzonase prior to immunoprecipitation and LC-MS/MS analysis. B) Efficient DNA digestion by Benzonase. Bioanalyzer traces show chromatin DNA fragments from the nuclear lysate before (–) and after (+) treatment with Benzonase. C-E) Volcano plots showing protein enrichment in Zld and Fib IPs. Data are presented as –log10 p-value vs. log2(IP/Input). IPs were performed using an anti-Zld antibody under standard conditions (C), with stringent 2M urea washes (D), or using an anti-Fib antibody as a control (E). Enriched proteins (FDR < 0.1) are highlighted in red. The FDR was determined using a combination of p-values and enrichment ratios in Perseus [61]. Key factors associated with chromatin modulation and transcription (C, D) or rRNA processing (E) are labeled. F) Overlap of enriched factors. Venn diagram showing the intersection of protein factors identified across the three IP experiments.

To define the protein complexes associated with Zld, we utilized a Zld antibody previously validated for high efficiency in ChIP-seq analysis [24]. This antibody exhibits high specificity; immunostaining analysis produced no signal in *zld-/-* germline clone embryos, in contrast to the robust signal observed in wild-type (WT) embryos. As a positive control for our IP-MS pipeline, we performed IPs using an antibody against Fibrillarin (Fib), a nucleolar protein involved in rRNA processing with a well-characterized set of associated factors [62]. Additionally, we conducted parallel analyses using anti-GFP antibodies in embryos endogenously expressing GFP-Zld.

### General characteristics of the enriched protein factors

Immunoprecipitation using the anti-Zld antibody was performed following both a standard protocol and a modified version incorporating stringent 2M urea wash steps to identify stable interactors. Under these conditions, we identified 138 and 152 enriched proteins, respectively (FDR < 0.1; Fig. 1, Supplementary Table 1). The two datasets showed a high degree of overlap, with 109 factors in common. In contrast, the control Fib-IP enriched 125 factors; of these, only 53 and 62 overlapped with the two anti-Zld IP-MS experiments, respectively.

Gene Ontology (GO) analysis revealed that Zld-associated factors were primarily related to RNA polymerase II (RNAPII) transcription, chromatin remodeling, and the regulation of chromatin organization (Table 1). Conversely, Fib-enriched factors were associated with rRNA modification and snoRNP complexes. We further validated our pipeline by cross-referencing Fib-enriched factors with the STRING database [63]; seven of the top ten predicted Fib interactors were significantly enriched (Fig. 1—figure supplement 1), while the remaining three were not detected in our MS dataset. Collectively, our GO and STRING analyses demonstrate that the Zld and Fib interactomes are distinct and align with their respective biological roles.

**Table 1.**
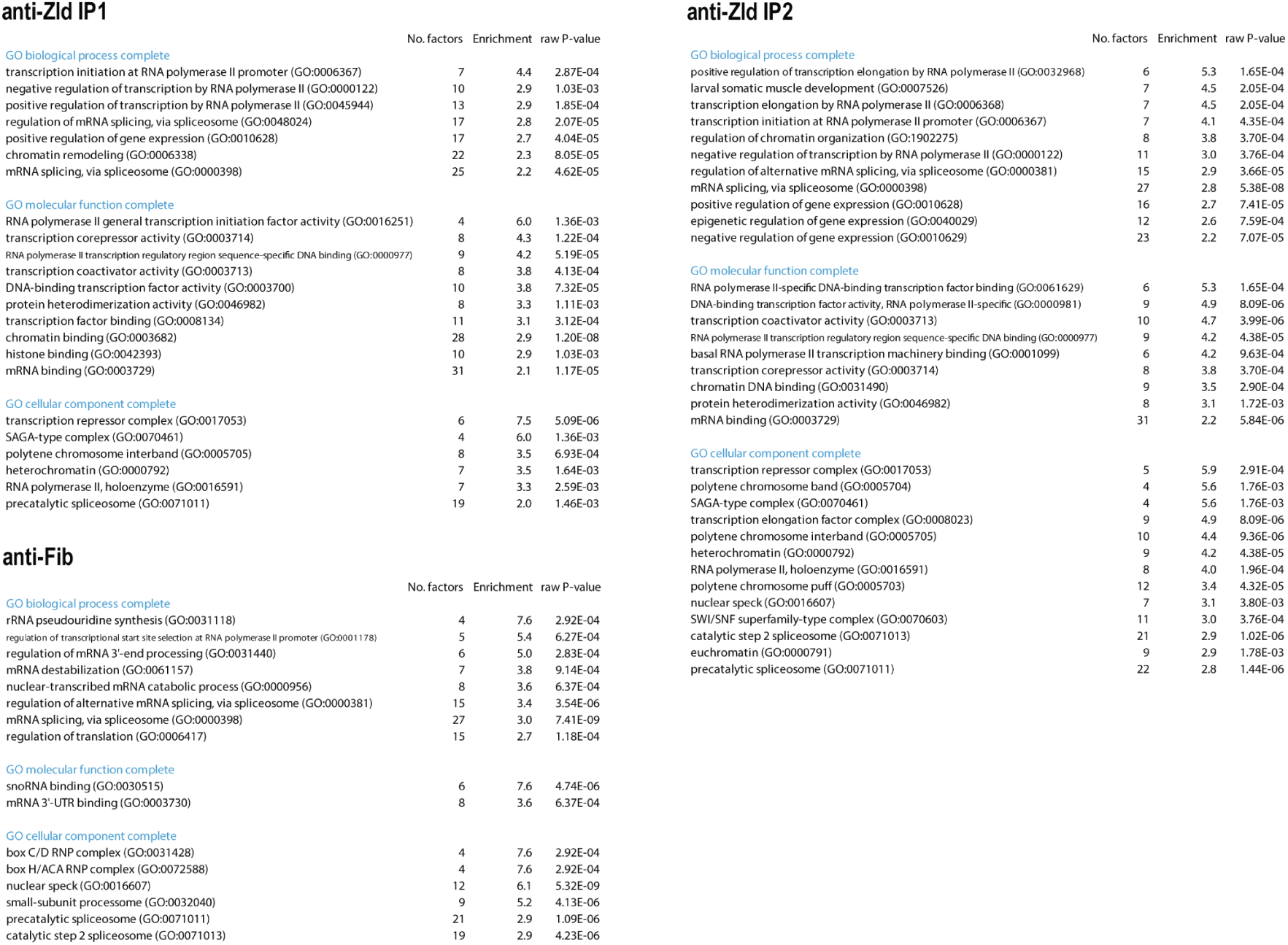
Top 10 enriched GO terms for the protein factors enriched in the anti-Zld IP-MS and anti-Fib IP-MS.

**Table 2.** Top 10 enriched GO terms for the proteins enriched in anti-GFP IP-MS.

| anti-GFP(#1) IP1 |  |  |  | anti-GFP(#2) |  |  |  |
| --- | --- | --- | --- | --- | --- | --- | --- |
|  | No. factors | Enrichment | raw P-value |  | No. factors | Enrichment | raw P-value |
| **GO biological process complete** |  |  |  | **GO biological process complete** |  |  |  |
| ATP-dependent chromatin remodeling (GO:0043044) | 7 | 14.7 | 1.38E-06 | ecdysone receptor-mediated signaling pathway (GO:0035076) | 3 | 47.9 | 1.11E-04 |
| regulation of chromatin assembly or disassembly (GO:0001672) | 4 | 13.5 | 4.11E-04 | nucleosome mobilization (GO:0042766) | 4 | 28.4 | 3.00E-05 |
| regulation of neuroblast proliferation (GO:1902692) | 5 | 11.6 | 1.34E-04 | regulation of chromatin assembly or disassembly (GO:0001672) | 4 | 19.7 | 9.58E-05 |
| nucleosome organization (GO:0034728) | 8 | 11.0 | 1.41E-06 | regulation of neuroblast proliferation (GO:1902692) | 5 | 16.8 | 2.15E-05 |
| chromatin assembly or disassembly (GO:0006333) | 5 | 8.8 | 4.07E-04 | ATP-dependent chromatin remodeling (GO:0043044) | 5 | 15.2 | 3.26E-05 |
|  | 7 | 6.0 | 2.13E-04 | positive regulation of transcription by RNA polymerase II (GO:0045944) | 10 | 12.5 | 8.38E-09 |
| DNA repair (GO:0006281) | 7 | 5.3 | 4.38E-04 | negative regulation of neurogenesis (GO:0050768) | 4 | 12.2 | 4.67E-04 |
| negative regulation of transcription, DNA-templated (GO:0045892) | 8 | 5.0 | 2.51E-04 | chromatin assembly or disassembly (GO:0006333) | 4 | 10.2 | 8.39E-04 |
|  |  |  |  | regulation of defense response (GO:0031347) | 5 | 8.9 | 3.16E-04 |
| **GO molecular function complete** |  |  |  | regulation of immune response (GO:0050776) | 5 | 8.0 | 4.93E-04 |
| nucleosome binding (GO:0031491) | 5 | 18.3 | 2.19E-05 | regulation of response to biotic stimulus (GO:0002831) | 5 | 7.3 | 7.36E-04 |
| DNA helicase activity (GO:0003678) | 4 | 14.7 | 3.19E-04 | regulation of transcription, DNA-templated (GO:0006355) | 19 | 5.4 | 7.00E-11 |
| transcription corepressor activity (GO:0003714) | 5 | 14.7 | 5.23E-05 |  |  |  |  |
| DNA-binding transcription factor activity, RNA polymerase II-specific (GO:0000981) | 4 | 11.7 | 6.47E-04 | **GO molecular function complete** |  |  |  |
| transcription factor binding (GO:0008134) | 6 | 6.4 | 4.64E-04 | nucleosome binding (GO:0031491) | 5 | 26.6 | 3.38E-06 |
| double-stranded DNA binding (GO:0003690) | 8 | 6.3 | 5.46E-05 | chromatin DNA binding (GO:0031490) | 4 | 13.5 | 3.34E-04 |
| ATP binding (GO:0005524) | 13 | 2.7 | 6.89E-04 | histone binding (GO:0042393) | 5 | 10.0 | 1.92E-04 |
|  |  |  |  | double-stranded DNA binding (GO:0003690) | 6 | 6.8 | 2.71E-04 |
| **GO cellular component complete** |  |  |  | transcription regulator activity (GO:0140110) | 7 | 6.3 | 1.25E-04 |
| SWI/SNF complex (GO:0016514) | 3 | 18.9 | 1.10E-03 | sequence-specific DNA binding (GO:0043565) | 6 | 6.2 | 4.52E-04 |
| histone deacetylase complex (GO:0000118) | 5 | 16.9 | 2.99E-05 |  |  |  |  |
| brahma complex (GO:0035060) | 4 | 16.0 | 2.43E-04 | **GO cellular component complete** |  |  |  |
|  |  |  |  | FACT complex (GO:0035101) | 2 | 63.9 | 1.35E-03 |
| **anti-GFP(#1) IP2** |  |  |  | ACF complex (GO:0016590) | 2 | 63.9 | 1.35E-03 |
|  |  |  |  | nuclear euchromatin (GO:0005719) | 3 | 27.4 | 3.70E-04 |
| **GO biological process complete** |  |  |  | SWI/SNF complex (GO:0016514) | 3 | 27.4 | 3.70E-04 |
| nuclear cell cycle DNA replication initiation (GO:1902315) | 3 | 40.1 | 1.89E-04 | RSC-type complex (GO:0016586) | 3 | 21.3 | 6.65E-04 |
| regulation of chromatin assembly or disassembly (GO:0001672) | 4 | 16.5 | 1.93E-04 | histone deacetylase complex (GO:0000118) | 4 | 19.7 | 9.58E-05 |
| protein-DNA complex subunit organization (GO:0071824) | 7 | 8.1 | 3.24E-05 | transcription repressor complex (GO:0017053) | 3 | 17.4 | 1.08E-03 |
| DNA conformation change (GO:0071103) | 7 | 6.2 | 1.52E-04 | brahma complex (GO:0035060) | 3 | 17.4 | 1.08E-03 |
| regulation of transcription by RNA polymerase II (GO:0006357) | 11 | 4.5 | 2.34E-05 | polytene chromosome (GO:0005700) | 9 | 7.2 | 3.93E-06 |
| negative regulation of cellular metabolic process (GO:0031324) | 11 | 3.6 | 1.77E-04 |  |  |  |  |
|  |  |  |  | **GO molecular function complete** |  |  |  |
| sequence-specific double-stranded DNA binding (GO:1990837) | 6 | 7.0 | 2.76E-04 |  |  |  |  |
|  |  |  |  | **GO cellular component complete** |  |  |  |
| protein-DNA complex (GO:0032993) | 5 | 9.9 | 2.21E-04 |  |  |  |  |
| nuclear chromatin (GO:0000790) | 9 | 6.7 | 8.65E-06 |  |  |  |  |
| polytene chromosome (GO:0005700) | 7 | 4.7 | 7.81E-04 |  |  |  |  |

### Functional categories of the Zld interactome

To generate a consolidated Zld interactome, we integrated results from both anti-Zld IP experiments, excluding factors with less than a two-fold enrichment (log_2_(IP/Input) < 1) in either dataset. This yielded 155 factors, 93 of which were specific to the Zld IPs (Fig. 2A).

**Fig. 2.**
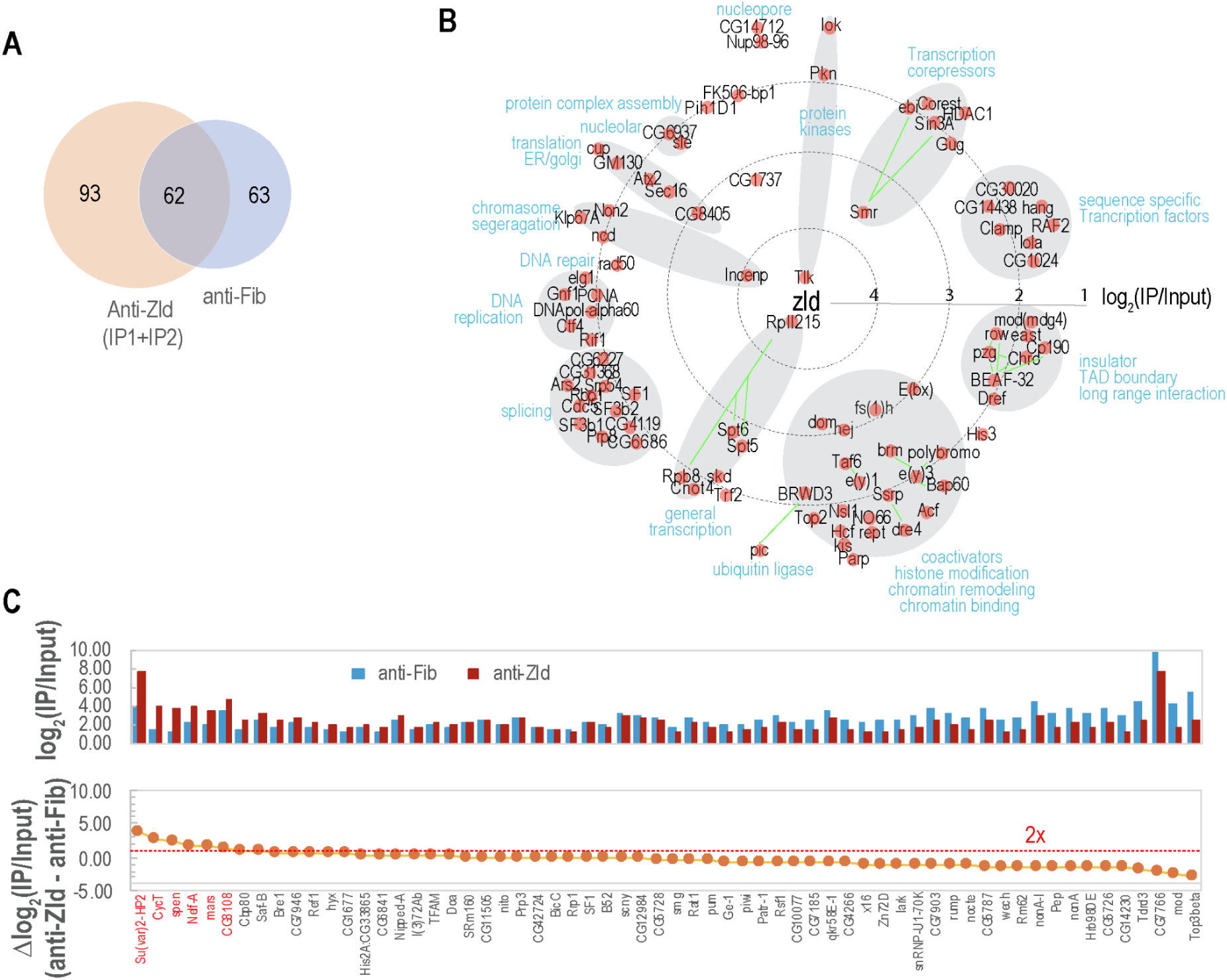
Quantitative and functional analysis of the Zelda (Zld) interactome. A) Overlap of protein factors enriched in anti-Zld and anti-Fib IPs. A total of 155 factors met the dual criteria of an FDR < 0.1 in at least one anti-Zld experiment and a minimum two-fold enrichment (log2(IP/Input)) ≥ 1) in the other. in the other. Of these, 62 factors were also enriched in the anti-Fib control, resulting in 93 Zld-specific interactors. B) Interaction map of Zld-specific factors. Factors specific to the anti-Zld experiments are grouped by functional relatedness. The distance of each factor from the center is inversely proportional to its average enrichment in the two anti-Zld IP-MS datasets. Green lines indicate, in selected cases, subunits of the same complex or previously reported physical interactions. C) Relative enrichment of shared factors. Differential enrichment for proteins identified in both anti-Zld and anti-Fib IPs. Data are plotted as the difference in log2(IP/Input) values between the two IPs. The red horizontal line indicates a two-fold enrichment difference (Δlog2(IP/Input) = 1).

As shown in the interaction map (Fig. 2B), the most prominent group of interactors consists of factors involved in histone acetylation and nucleosome remodeling. These include the coactivators dCBP and Fsh, as well as subunits of major nucleosome remodeling complexes [64,65], such as the NURF complex (E(bx)), the PBAP/BAP complexes (Brm, SAYP, Polybromo(PB), and Bap60), and the ACF complex. We also observed strong enrichment of Domino (DOM), a subunit of the TIP60 and SWR1 complexes [66], and the histone chaperone FACT complex (Dre4 and Ssrp) [67]. The recruitment of such a diverse array of remodelers suggests that Zld may directly recruit these machines to establish accessibility at enhancers, reminiscent of the pioneer factor GAF [68,69].

While general transcription factors (GTFs) were sparsely represented, we observed striking enrichment of the RNA polymerase II large subunit (Rpb1; hereafter referred to as RNAPII), which ranked as the second-highest Zld-specific interactor. Several of its associated factors were also enriched, including the negative elongation factor (NELF-A), the elongation factor P-TEFb (CycT), Spt5, Spt6(Fig. 2B-C). NELF-A and CycT were also enriched in the anti-Fib IP, but at much lower levels than observed in the Zld IPs. Beyond these factors required for chromatin accessibility and transactivation—the coactivators (e.g., dCBP and Fsh), nucleosome remodelers, and the RNAPII complex—several factors involved in transcriptional repression were also enriched. Among the most highly enriched proteins was the corepressor Smrter (Smr), identified along with its established partners Ebi and Sin3A [55,56]. This finding is particularly noteworthy as Zld has been shown to contain a transcription repression domain [50], suggesting that Zld may recruit Smr to modulate or “buffer” the output of its target genes.

Most intriguingly, the single most enriched factor in our anti-Zld IPs, surpassing even RNAPII, was the kinase Tlk. Tlk is an evolutionarily conserved protein whose activity is tightly coupled to the cell cycle, peaking during S-phase to regulate chromatin assembly [57,70]. Given the rapid, synchronous nuclear divisions characteristic of early embryo development, Tlk is a prime candidate for coordinating Zld-mediated ZGA with the embryonic mitotic cycle.

We also detected several factors associated with chromatin insulators and the maintenance of Topologically Associating Domain (TAD) boundaries, including BEAF-32, CP190, Pzg, and Row. While the enrichment for these factors was moderate, their presence is supported by several genomic observations. Zld is required for the *de novo* formation of TAD boundaries in the early embryo, and its binding motifs are significantly enriched at these sites [71,72]. Furthermore, the BEAF-32 motif was identified as a top-ranking enriched sequence within Zld ChIP-seq peaks in both larval neuroblasts and early embryos [73]. Collectively, our data provide a biochemical link between Zld and the spatial organization of higher-order chromatin structure during the MZT.

Other enriched factors further underscore Zld’s role as a cooperative regulator. For example, the transcription factor Clamp, which cooperatively binds and activates enhancers with Zld

[73–75], was also identified. Additionally, the weak enrichment of DNA replication factors may be explained by the widespread stalling of DNA replication at transcriptionally engaged, Zld-activated genes [76]. We also noted the enrichment of CG1737, a factor of unknown function, which was also previously identified in pull-downs for Chro [77] and HP1 [78]. However, whether this factor is functionally related to Zld requires further investigation.

In summary, our Zld interactome comprises a diverse repertoire of factors, suggesting that Zld does not merely “open” chromatin but acts as a central scaffold that bridges the maternal-to-zygotic transition by integrating transcriptional activation with the unique temporal demands of early embryogenesis.

### Validation of the Zld interactome using endogenously tagged GFP-Zld

To corroborate our findings, we performed IP-MS using a fly line expressing endogenously GFP-tagged Zld [37]. Across three biological replicates, we identified 52 high-confidence interactors (Fig. 3), 32 of which overlapped with our primary anti-Zld IP-MS results. Consistent with our initial findings, the GFP-Zld IPs were significantly enriched for subunits of the NURF, FACT, and BAP/PBAP complexes, as well as TAD boundary factors including Row, Pzg, and Chro. Notably, we observed strong enrichment for the mismatch repair factor Msh6 across GFP replicates, even though it had not reached statistical significance in the initial anti-Zld IP-MS. Msh6 has been shown to be associated with replication factories [79], and its enrichment here - like that of other DNA replication factors - may reflect the stalling of replication at Zld-dependent transcription sites, though a more direct role in Zld function cannot be excluded.

**Fig. 3.**
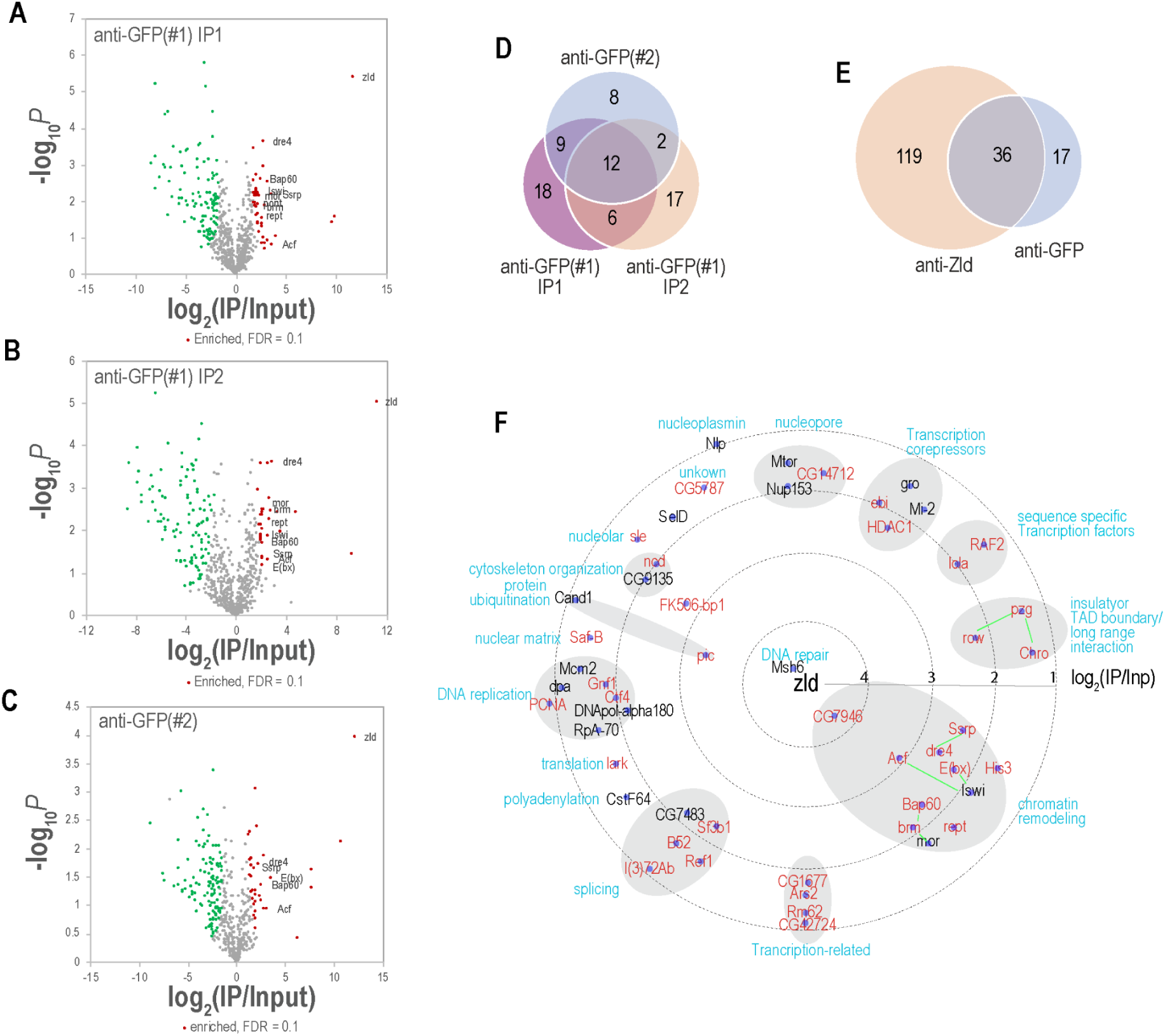
IP–MS analysis using anti-GFP antibodies in embryos expressing endogenously tagged GFP-Zld. Immunoprecipitation was performed using two distinct anti-GFP antibodies: anti-GFP(#1) and anti-GFP(#2). For anti-GFP(#1), experiments were conducted under two different washing conditions to assess interaction stability. Protein enrichment was analyzed as described in Figure 1. A) Enrichment of protein factors using anti-GFP (#1) under standard conditions. Volcano plot showing-log10P vs. log2(IP/Input). Factors enriched at an FDR < 0.1 are highlighted in red; select interactors associated with chromatin structure are labeled. B) Enrichment using anti-GFP (#1) under high-stringency conditions. Similar to (A), but including a 2M urea wash step during the immunoprecipitation. C) Enrichment using anti-GFP (#2). Similar to (A), but utilizing a different anti-GFP antibody. D) Overlap of GFP-Zld interactors across experiments. Venn diagram showing the distribution of enriched factors across the three anti-GFP IP-MS experiments. E) Comparison of anti-Zld and anti-GFP interactomes. The factors from the three anti-GFP experiments were consolidated and compared to the total list of factors enriched in the anti-Zld IP-MS experiments. F) Functional interaction map of GFP-Zld-associated factors. Factors enriched in the anti-GFP IP-MS experiments are arranged by functional relatedness, with distance from the center inversely proportional to average enrichment. Factors that were also identified in the anti-Zld experiments are highlighted in red.

The total number of enriched factors in the GFP-Zld IPs was notably smaller than in the anti-Zld IPs. We determined that this reduced coverage was primarily due to the proteolytic truncation of the GFP-Zld protein in the embryo. Immunostaining analysis revealed that N-terminal fragments (detected by anti-GFP but not by a C-terminal anti-Zld antibody) were located predominantly in the apical cytoplasmic layer (Fig. 3—figure supplement 1). Within the nuclei, both anti-GFP and anti-Zld signals were present; however, quantitative image analysis showed that the spatial overlap between these clusters was significantly lower than the overlap observed between clusters generated by two different fluorophore-conjugated secondary antibodies following anti-Zld primary antibody incubation in the same embryos. This indicates that while full-length Zld is nuclear, these truncated fragments localize to the nucleus independently. This protein fragmentation was confirmed by western blot and MS peptide mapping, which showed a sharp decline in spectral counts C-terminal to amino acid 1023 (Fig. 3—figure supplement 2). This intrinsic instability suggests that while endogenous tagging is a powerful tool, it can impact the efficiency of interactome discovery and must be accounted for in both biochemical and live-imaging analyses.

### Genomic profiling confirms Zld co-localization with key factors

To confirm these biochemical associations in vivo, we performed CUT&RUN for a subset of the most highly enriched factors, including dCBP, Fsh, Bap, Brm, Ssrp, Smr, and Tlk. We found that all these factors co-localized extensively at Zld binding sites (Fig. 4).

**Figure 4.**
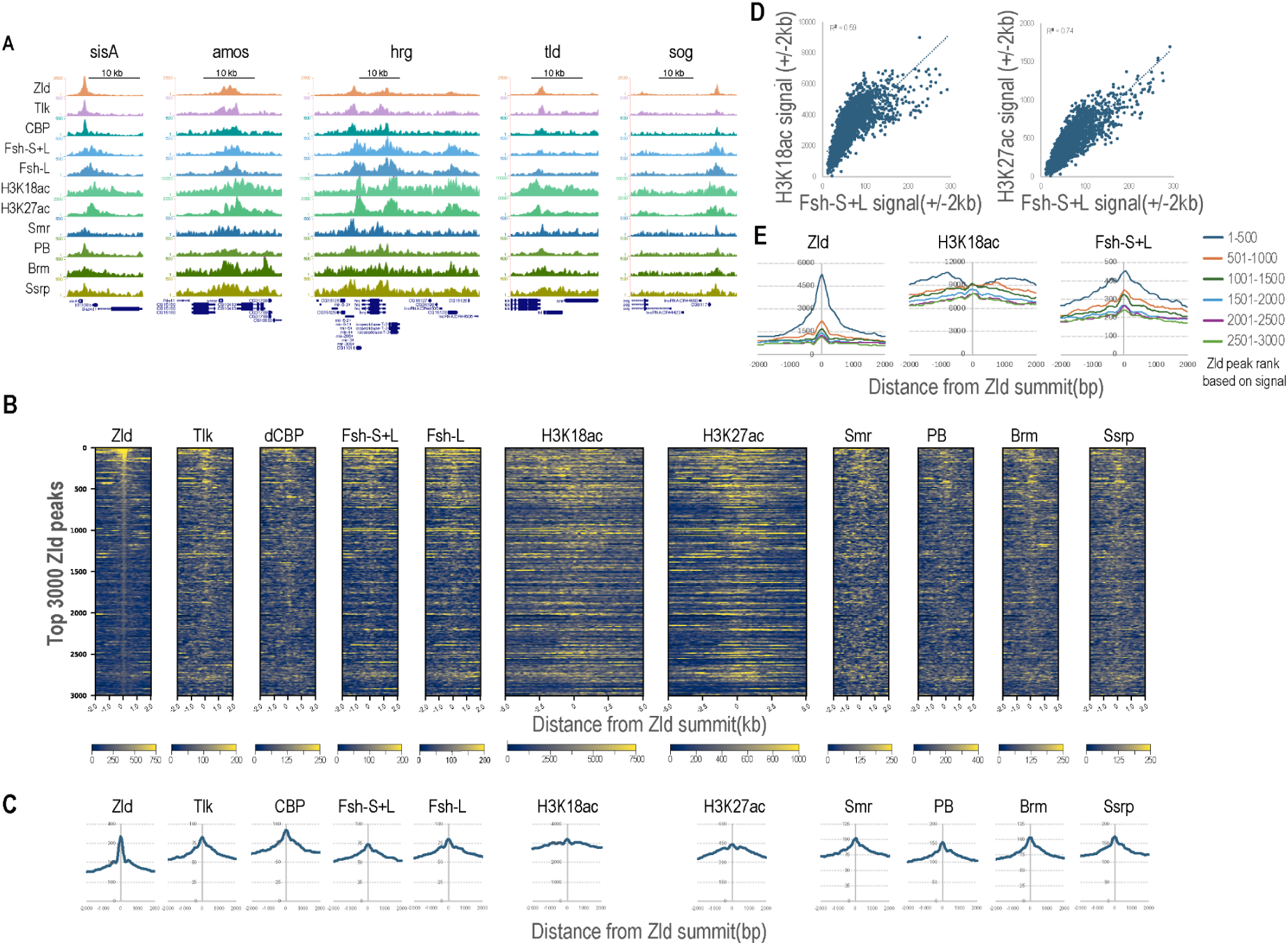
Genomic validation of Zld-associated factors via CUT&RUN. A) Representative genomic tracks of Zld-associated factors. CUT&RUN signal profiles for the indicated antibodies around the sisA, amos, hrg, tld, and sog loci. The experiments were performed in formaldehyde-fixed cycle 12 embryos. B) Global occupancy of Zld interactors at Zld-bound sites. Heatmaps displaying the CUT&RUN signal for the indicated factors centered on the top 3,000 Zld peaks. Peaks are sorted by fold enrichment as determined by MACS.. C) Average occupancy profiles at Zld peaks. Composite signal profiles for each antibody centered on all identified Zld peaks. D) Correlation of Fsh occupancy with histone acetylation. Correlation of Fsh (Fsh-S + Fsh-L) signal with H3K18ac and H3K27ac enrichment around Zld-bound regions. The result for Fsh-L is very similar (not shown). E) Fsh recruitment correlates with Zld peak intensity despite local histone depletion. Fsh recruitment correlates with Zld peak intensity despite local histone depletion. Zld peaks were ranked by fold enrichment and divided into cohorts of 500. Average signal profiles for each cohort demonstrate that at high-intensity Zld peaks, Fsh signal remains robust directly over the peak center, even where H3K18ac levels are significantly lower than in flanking sequences. Results for Fsh-L were highly similar (data not shown). The experiments were performed in formaldehyde-fixed cycle 13 embryos.

Fsh is expressed as two variants—a short (Fsh-S) and a long (Fsh-L) isoform—resulting from alternative splicing. We utilized two distinct Fsh antibodies: one specific to the long isoform (ID173) and another recognizing both isoforms (ID166) [80]. Both antibodies detected nearly identical binding patterns around Zld sites (Fig. 4—figure supplement B), suggesting that Fsh-L may be the predominant isoform at these regions. In support of this, our anti-Zld IP-MS analysis showed that the enrichment of peptides unique to the long isoform was approximately two-fold higher than the enrichment calculated from peptides shared by both isoforms (Fig. 4—figure supplement 1C–E). Given that these two isoforms are reported to be present at similar levels in the cell [80], this discrepancy in enrichment suggests that Fsh-L is the primary isoform recruited to Zld-bound regions, although direct evidence would be required to confirm this definitively.

Consistent with previous ChIP-seq results [27], we observed robust H3K18ac and H3K27ac signals flanking these Zld-occupied regions; notably, the signals for these two marks were highly correlated (Fig. 4—figure supplement 1F). Fsh occupancy generally correlated with broad domains of H3K18ac and H3K27ac across the genome, consistent with its known function in promoting transcriptional activation by binding to acetylated histones [54,81,82]. Notably, however, a distinct pattern emerged at the centers of strong Zld peaks. Despite the local depletion of acetylated histones directly over these sites—likely due to nucleosome displacement—focal Fsh signals remained highly concentrated, often exceeding the intensity of the flanking acetylated regions (Fig. 4D–E). This pattern became significantly more pronounced in cycle 13 than in cycle 12. These observations suggest that while Fsh may associate with the genome via histone acetylation globally, its concentrated recruitment to these specific centers is likely mediated by a distinct mechanism, such as direct physical interaction with Zld or other enhancer-associated factors.

### Recruitment of RNAPII to distal Zld binding sites

To investigate the striking RNAPII enrichment observed in our IP-MS, we performed RNAPII CUT&RUN across early embryonic stages. Surprisingly, we found that the RNAPII signal was not restricted to promoters; instead, it was broadly observed across nearly all Zld-bound regions, including distal intergenic peaks (Fig. 5A–C). Notably, when comparing RNAPII enrichment at these intergenic Zld peaks to their nearest potential target promoters, the signal at the distal Zld sites was often significantly more robust (Fig. 5D). This pattern was consistent at active enhancers for genes such as *Kr* and *sog*, as well as at distal enhancers for genes like *soxN*, which are Zld-bound but not yet actively transcribed. This widespread distal localization contrasts with previous ChIP-chip [83] and ChIP-seq [84] studies, which primarily localized RNAPII to transcription start sites(TSS).

**Figure 5.**
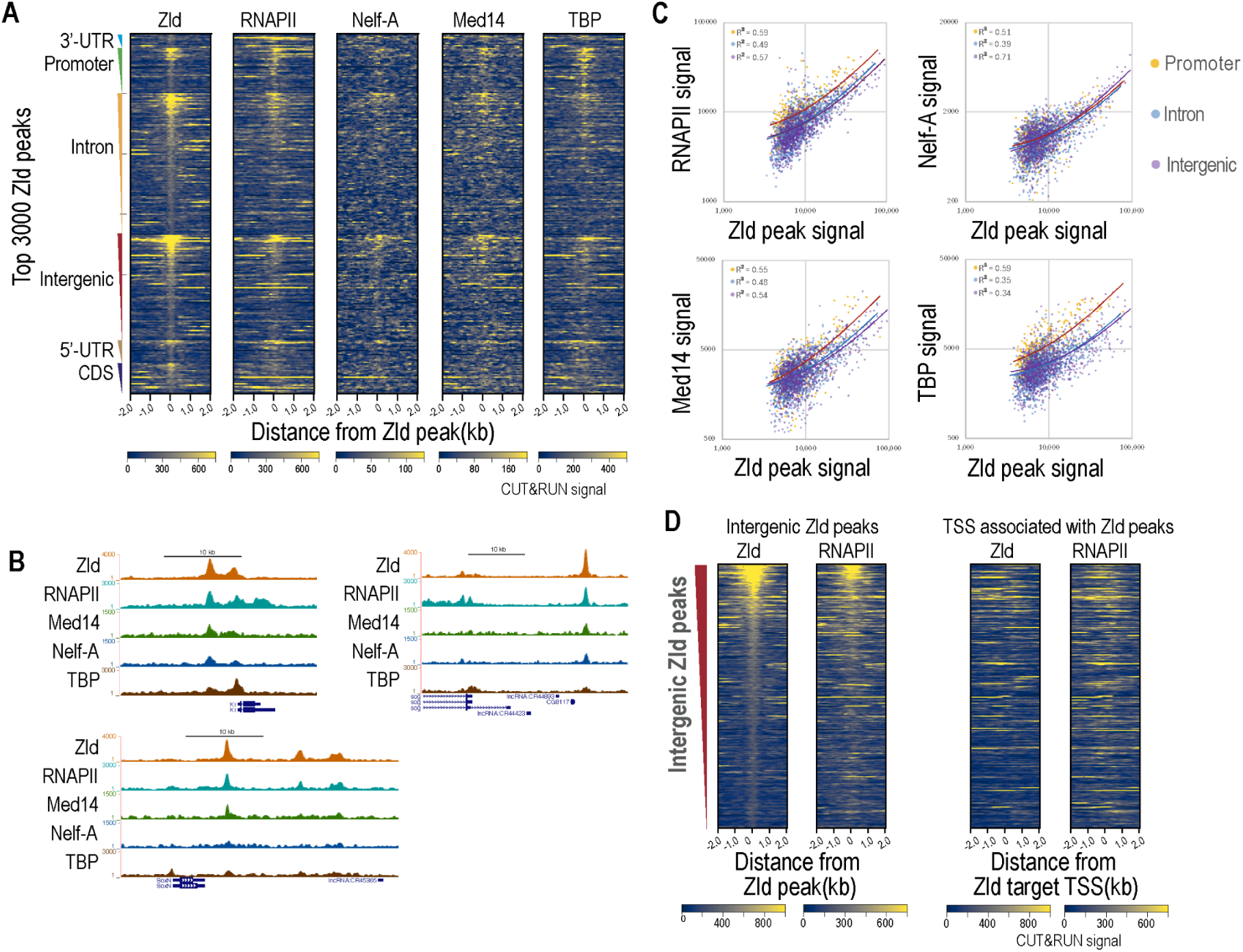
Zld recruits RNAPII, NELF-A, Med14, and TBP to distal and proximal binding sites. A) Global occupancy of transcription machinery across genomic features. Heatmaps displaying CUT&RUN signals for the indicated factors. Zld peaks are categorized by genomic location (3’ UTR, intron, intergenic, CDS, and 5’ UTR) and sorted within each category by Zld fold enrichment. B) Representative genomic tracks. Signal profiles for Zld, RNAPII, NELF-A, Med14, and TBP around selected gene loci, demonstrating co-localization at both promoters and distal enhancers. C) Correlation of factor occupancy with Zld binding. Scatter plots and correlation coefficients showing the relationship between Zld signal and each identified factor within a ±250 bp window around Zld peaks. While all factors correlate with Zld, TBP displays a relatively higher signal bias toward promoters. D) Comparison of RNAPII occupancy at intergenic enhancers versus target promoters. Heatmaps centered on intergenic Zld peaks (left) and their corresponding nearest transcription start sites (TSS; right). RNAPII signals are generally more robust at the intergenic Zld-bound enhancers than at the potential target promoters.

RNAPII is not alone in its association with these distal regions. As shown in Fig. 5, the RNAPII-associated factor NELF-A, which was also identified in our IP-MS, as well as the Mediator subunit Med14 and the general transcription factor (GTF) TBP, all co-localized with Zld at distal enhancers. Collectively, these CUT&RUN results suggest that Zld may serve as a scaffold to recruit RNAPII, the NELF complex, the Mediator complex, and, to a lesser extent, TBP, directly to its binding sites. This distal assembly of the transcriptional machinery likely facilitates the rapid transactivation of Zld target genes by pre-configuring the transcriptional hub prior to or concurrent with promoter communication.

### High-resolution imaging for Zld-RNAPII colocalization

To visualize the spatial relationship between Zld and the transcriptional machinery, we performed immunostaining in early embryos using the Zeiss Airyscan detector system for super-resolution imaging. We observed that Zld and RNAPII form distinct sub-nuclear clusters (Fig. 6A, Fig. 6—figure supplement 1), consistent with the specialized transcription “hubs” previously described [35–37,85,86]. Both Zld and RNAPII clusters are dynamic during the mitotic cycle, with Zld clusters forming early and RNAPII clusters forming with a delay [87](Fig. 6—figure supplement 1)

**Fig. 6.**
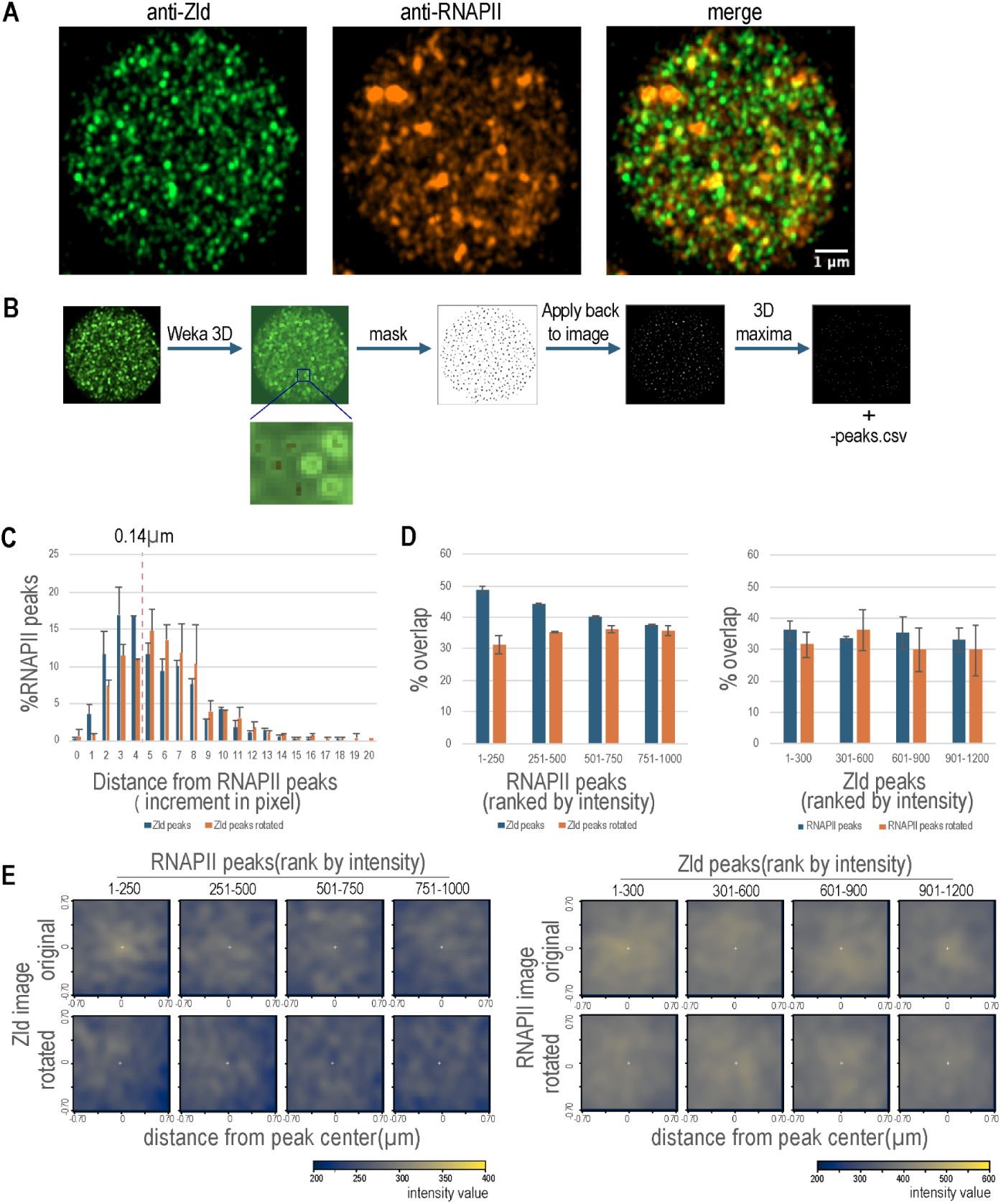
Colocalization of Zld and RNAPII in embryo. The embryos were stained with anti-Zld and anti-RNAPII (4H8) antibodies; high resolution images were acquired using the Zeiss Airyscan detection system. A) Visualizing sub-nuclear clusters. MIP images of 5 Z-slices from a mid-cycle 12 embryo. B) Image analysis workflow. Schematic illustrating the peak identification and cluster segmentation process performed in Fiji. C) Object-based 3D distance analysis. Frequency distribution of 3D Euclidean distances from the top 250 RNAPII peaks to the nearest Zld peak. Distances were compared against a 90° rotated control to establish a stochastic noise floor. A significantly higher percentage of RNAPII peaks are within 0.14 µm (4 pixels) of a Zld peak compared to the rotated control. D) Enrichment of brightest RNAPII hubs. Percentage of peak overlap at a 0.14 µm cutoff for intensity-ranked peak cohorts. The most significant colocalization occurs within the top 250 brightest RNAPII spots (∼49% vs. ∼31% in rotated control; n = 2 nuclei). E) Signal intensity heatmaps. Average RNAPII signal centered around Zld peaks and Zld signal centered around RNAPII peaks. For comparison, heatmaps generated after rotating target images by 90° are shown. Significant Zld enrichment is observed at the most intense RNAPII peaks.

To rigorously quantify Zld–RNAPII association and account for potential chance overlaps in the crowded nuclear environment, we employed object-based 3D distance analysis compared against a rotated control. In this control, one channel and its corresponding nuclear mask were rotated 90° relative to the other to randomize the spatial relationship while maintaining original cluster density and morphology. In mid-cycle 12 embryos, we found that the brightest RNAPII clusters showed a significant preferential association with Zld. Specifically, 48.9 ± 1.0% of the top 250 RNAPII spots were located within 0.14 µm of a Zld cluster, representing a ∼1.6-fold enrichment over the randomized baseline (31.2 ± 3.0%; Fig. 6D, left panel). This enrichment was intensity-dependent, with lower-ranked RNAPII cohorts approaching the stochastic overlap levels (Fig. 6D, left panel). This preferential association also diminished later in the mitotic cycle as RNAPII clusters became weaker (Fig. 6C–D, Fig. 6—figure supplement 1B–C).

As a secondary validation, we used heatmap analysis to examine signal distribution around peak centers (Fig. 6E, Fig. 6—figure supplement 1D). Consistent with our object-based findings, Zld signal was significantly enriched around the top 250 RNAPII peaks, a finding consistent with the functional dependence of these transcriptional hubs on Zld [88]. Conversely, RNAPII signal was only modestly enriched around the brightest Zld peaks. This asymmetry suggests that the Zld–RNAPII association is likely highly dynamic; furthermore, the high-intensity RNAPII clusters captured in our imaging analysis likely correspond to elongating polymerases at gene bodies, which may obscure the weaker RNAPII signals associated with Zld at enhancers. Taken together, these data demonstrate that Zld and the core transcriptional machinery are spatially coordinated, particularly at the most active transcriptional hubs.

### Zld clusters preferentially associate with target gene transcription loci

We next investigated whether these Zld hubs are specifically localized to Zld-target genes. Using smiFISH to detect nascent RNA transcripts, we measured the 3D distance between Zld protein clusters and active transcription foci, comparing Zld-target genes directly to non-target controls.

We found that transcription foci for the endogenous Zld-target gene *zen* were located significantly closer to Zld clusters than those of the non-target gene *dfd* (Fig.7A–D). This Zld-dependent spatial coordination was further confirmed using a *lacZ* reporter driven by the *hb* stripe enhancer [8]—while the wild-type reporter showed close proximity to Zld clusters, the mutated *lacZ* reporter lacking Zld-binding sites was located significantly further away (Fig. 7E–H). These results demonstrate that Zld clusters are functional transcriptional hubs recruited to specific genomic loci through direct Zld-DNA interactions, and that the loss of these binding sites results in the physical dissociation of the gene from the hub.

**Fig. 7.**
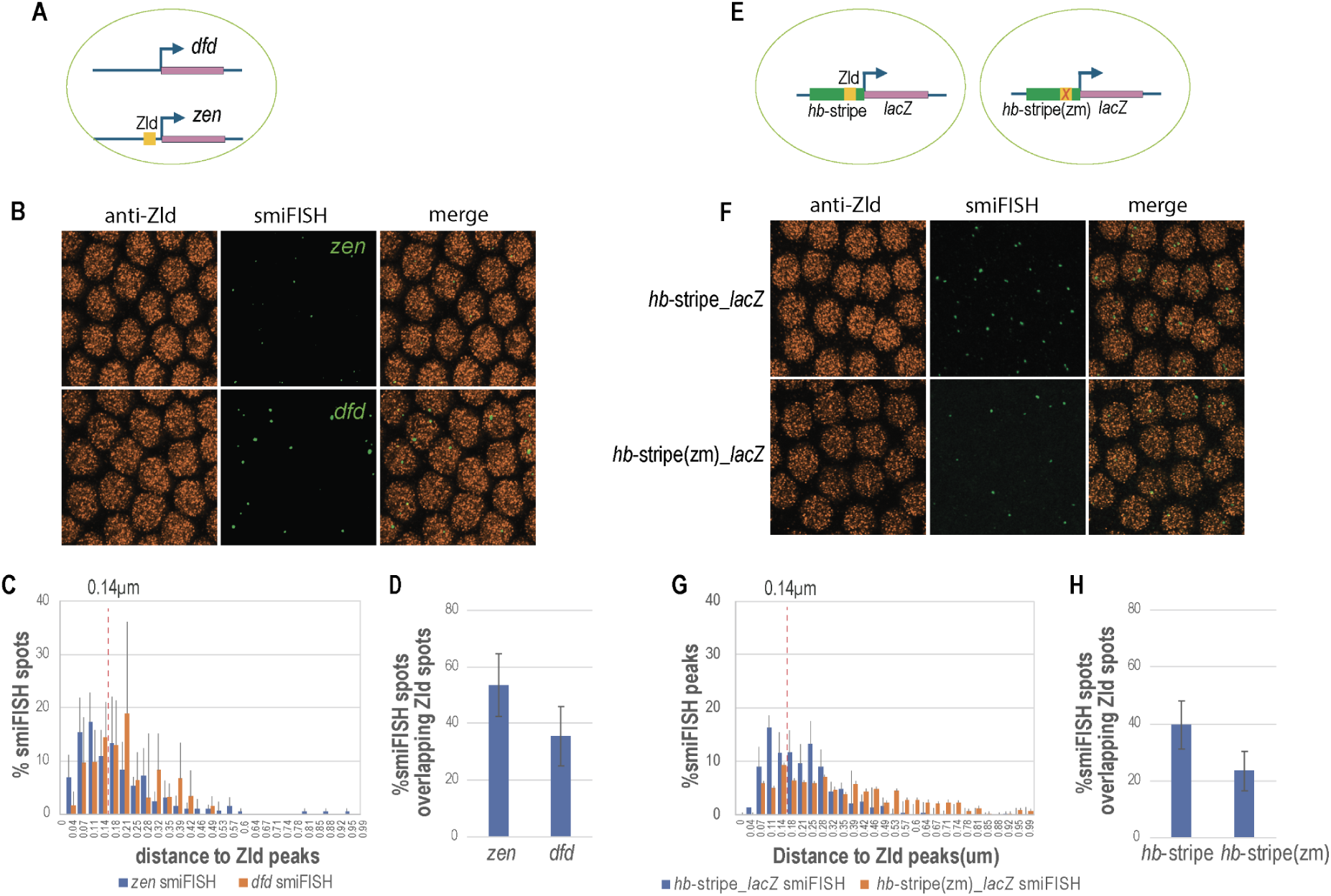
Co-localization of Zld clusters with active transcription foci of Zld target genes. High-resolution smiFISH and immunostaining using an anti-Zld antibody were performed in fixed embryos. The smiFISH method detects both individual mRNA molecules and high-intensity clusters (transcription foci) representing sites of nascent transcription. Zld peaks and transcription foci were identified using the 3D Maxima plugin in Fiji. Nearest-neighbor distances between transcription foci and their closest Zld peak were calculated as described in Figure 6. A) Experimental design for endogenous target validation. Schematic of zen (a Zld target) and dfd (a non-target). Probes for both genes were used simultaneously in the same embryos, allowing for intra-nuclear comparison. B) Visualization of transcription foci and Zld hubs. Representative regions (25 μm × 25 μm) are shown from original 100 μm × 100 μm MIP of an early c14 embryo. Images were thresholded to specifically highlight high-intensity smiFISH signals at active transcription sites. C) Spatial distribution of transcription foci relative to Zld. Frequency distribution of the distances between zen or dfd smiFISH spots and the nearest Zld cluster. D) Quantitative co-localization of endogenous foci. Percentage of zen and dfd transcription foci co-localizing with Zld clusters within a 0.14 µm center-to-center cutoff. E) Reporter constructs for Zld-dependency testing. Schematics of the lacZ reporter transgenes driven by either the wild-type hb stripe enhancer (hb-stripe) or a version with mutated Zld binding sites (hb-stripe(zm)). F) Visualization of reporter transcription. MIP images of 25 µm × 25µm showing lacZ smiFISH signals in cycle 14 embryos for both the wild-type and Zld-binding site mutant transgenes. G) Distance distribution of reporter foci. Distribution of distances between lacZ transcription foci and the closest Zld clusters for both transgenes. H) Distance distribution of reporter foci. Distribution of distances between lacZ transcription foci and the closest Zld clusters for both transgenes.

### dCBP mediates Zld-dependent histone H3K18 acetylation and Fsh cluster formation

Zld binding is known to correlate with several histone acetylation marks—including H3K18ac, H3K27ac, and H4K8ac [27]—that are deposited by dCBP [89,90]. Fsh/BRD4 orthologs, on the other hand, bind acetylated histones to facilitate enhancer function [52–54,81,82,91]. Based on these findings and the strong enrichment of Fsh and dCBP in the anti-Zld IP-MS, we hypothesized that these two factors play critical roles in Zld-dependent transactivation.

BRD4 has been shown to form nuclear puncta with liquid-like condensate properties at super-enhancers in various mammalian cell types [92–94]. To characterize the in vivo relationship between Zld, dCBP-mediated acetylation, and Fsh, we performed immunostaining for H3K18ac and Fsh in wild-type, *nej* (dCBP) RNAi-knockdown, and *zld* mutant embryos

High-resolution Airyscan imaging in wild-type embryos revealed a vast number of distinct H3K18ac and Fsh clusters, reminiscent of the BRD4 condensates observed in mammalian cells [92–94](Fig. 8A). Given the specificity of the anti-H3K18ac signal for chromatin, these clusters likely represent individual or small groups of co-localized active enhancers. Analysis across early mitotic cycles showed that weak H3K18ac clusters are detectable as early as cycle 8 and increase in both number and intensity through subsequent cycles (Fig. 8—figure supplement 1). This cumulative deposition suggests that chromatin opening is a progressive process, likely reflecting the transition of zygotic enhancers from a ‘primed’ to a ‘fully active’ state.

**Fig. 8.**
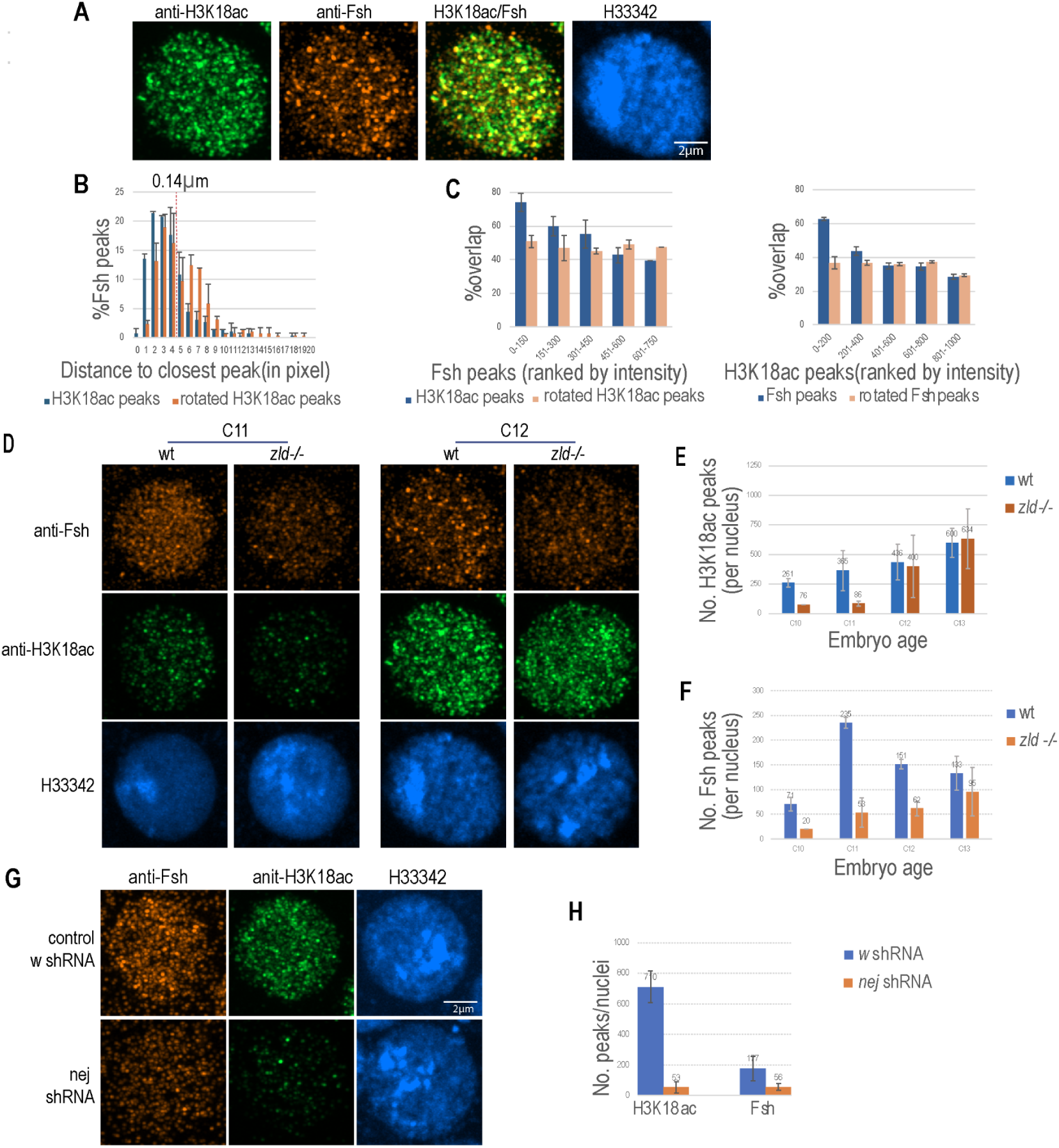
dCBP and Zld are required for H3K18ac and the formation of Fsh clusters. Immunostaining was performed in fixed embryos using anti-H3K18ac and anti-Fsh (ID166) antibodies. Images were acquired and analyzed as described in Figure 6. A) Representative sub-nuclear distribution. MIP images of a single nucleus from a cycle 12 embryo showing H3K18ac and Fsh clusters. B) Spatial correlation between Fsh and H3K18ac. Distribution of nearest-neighbor distances from the top 150 Fsh peaks to the closest H3K18ac peaks. The distribution obtained after rotating H3K18ac peak coordinates by 90° is shown as a control. C) Intensity-dependent colocalization. Percentage of Fsh peaks in each intensity-ranked cohort colocalizing with H3K18ac peaks (left), and vice versa (right), using a 0.14 µm center-to-center cutoff. D) Zld-dependency of Fsh and H3K18ac. MIP images from wild-type and zld–/– germline clone mutant embryos. The wild-type cycle 12 image is the same as shown in (A). E) Temporal dynamics of H3K18ac loss in zld mutants. Quantitative change in the number of H3K18ac clusters per nucleus in zld mutants compared to wild-type from cycles 10 to 13. F) Temporal dynamics of Fsh cluster loss in zld mutants. Quantitative change in the number of Fsh clusters per nucleus in zld mutants compared to wild-type from cycles 10 to 13. G) Loss of clusters upon dCBP knockdown. MIP images of embryos expressing shRNA targeting nej (encoding dCBP) or w (negative control). H) Quantification of dCBP-dependency. Comparison of the number of Fsh and H3K18ac clusters per nucleus in nej versus w RNAi embryos at cycle 13.

Consistent with Fsh recruitment by acetylated histones, object-based colocalization analysis revealed significant spatial overlap between Fsh and H3K18ac clusters, with the highest-intensity peaks exhibiting the strongest colocalization (Fig. 8B-C). This indicates a more stable association of Fsh with extensively acetylated, high-occupancy enhancers.

In *zld* mutant embryos (germline clones), both H3K18ac and Fsh signals were significantly reduced (Fig. 8D–F). The loss of H3K18ac was most pronounced during cycles 10–11, with a partial recovery by cycle 12. However, the brightest Fsh clusters remained absent, suggesting that while other factors may eventually contribute to acetylation, the assembly of large, Fsh-rich clusters is strictly Zld-dependent. Furthermore, RNAi knockdown of *nej* dramatically decreased both H3K18ac and Fsh clusters (Fig. 8G–H, Fig. 8—figure supplement 2), confirming that Fsh recruitment is dependent on dCBP-mediated acetylation.

### Fsh and dCBP are critical for the transactivation of Zld-dependent genes

To directly test the requirement of Fsh and dCBP in Zld-mediated transactivation, we analyzed the effects of RNAi knockdown on the transcription of Zld-dependent genes (*sala, tld,* and *hb*) and a Zld-independent control (*btd*) [23]. We performed smiFISH in embryos expressing shRNA targeting either *fs(1)h* or *nej*, using *white (w)* shRNA as a negative control. In our smiFISH analysis, individual mRNA molecules were detected as distinct puncta, while active transcription sites appeared as one or two high-intensity foci per nucleus. For *hb*, which has significant maternal RNA deposition, we quantified only the active transcription foci. For all other genes, we quantified the total number of both spot types across the Z-stack using the Fiji 3D Maxima plugin.

In addition to smiFISH, we performed immunostaining with an anti-RNAPII antibody to monitor transcriptional activity at the histone gene cluster, visualized as the Histone Locus Body (HLB), which is independent of Zld [88]. As shown in Fig. 9B–C, *fs(1)h* knockdown (Fig. 9—figure supplement 1) significantly reduced the transcription of *sala, tld,* and *hb*, but not that of the Zld-independent gene *btd*. Furthermore, the RNAPII signal at the HLB was only modestly reduced in *fs(1)h* RNAi embryos compared to controls. These results indicate that Fsh is specifically required for the transcription of Zld-dependent genes.

**Fig. 9.**
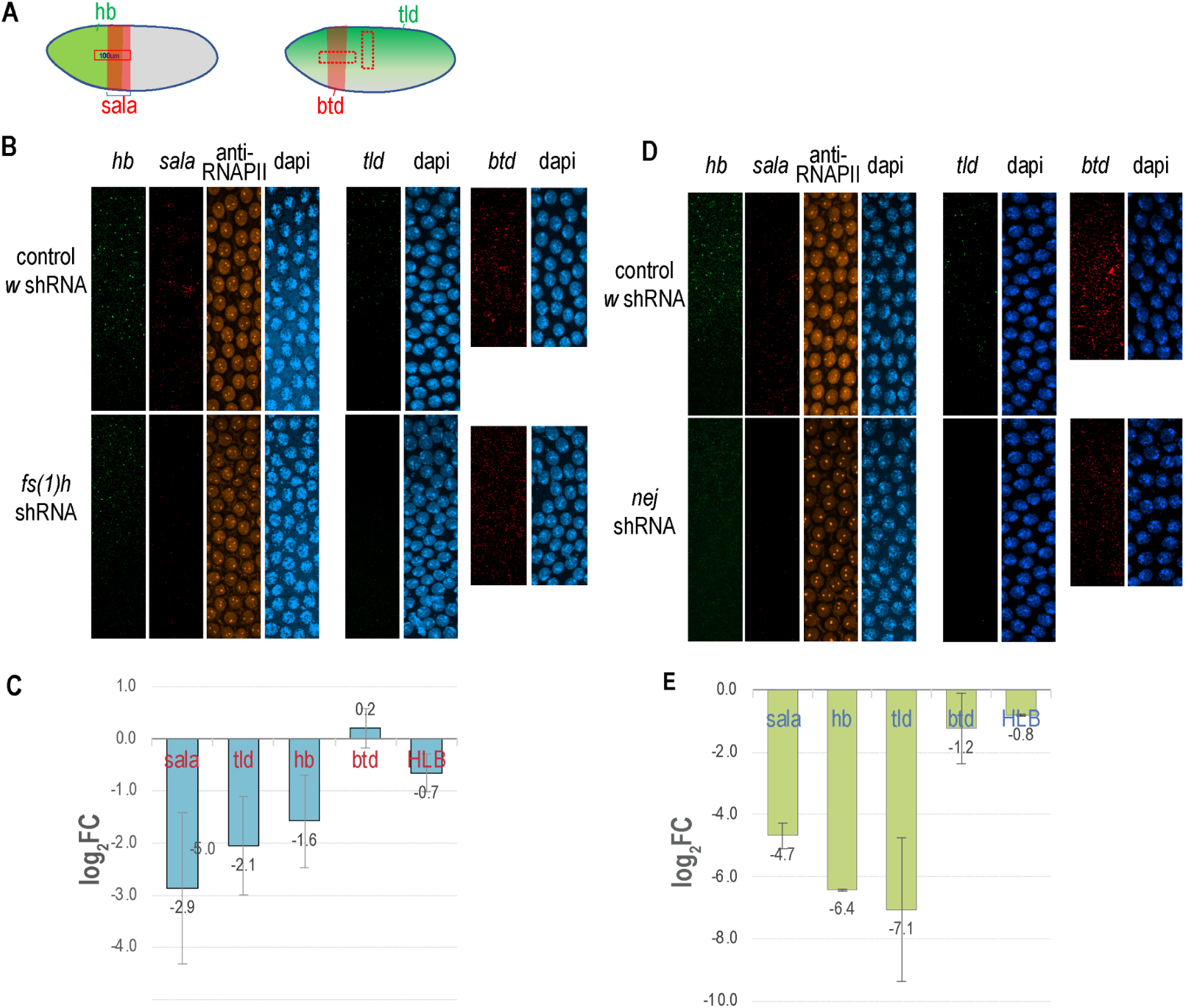
Requirement of Fsh and dCBP for the transactivation of Zld-dependent genes. smiFISH for sala, tld, hb, and btd was performed in embryos expressing shRNA targeting fs(1)h or nej (dCBP), with white (w) shRNA serving as a negative control. Following smiFISH hybridization, embryos were immunostained with an anti-RNAPII antibody. The number of individual mRNA molecules for sala, tld, and btd was quantified within the indicated embryonic regions using the Fiji 3D Maxima plugin. For hb, the number and intensity of bright nuclear foci (representing active transcription) were determined, and the integrated intensity (sum of focus intensities) was calculated for each scanned area. Additionally, the average intensity of RNAPII signal at the Histone Locus Body (HLB) was measured per embryo. A) Spatial domains of analyzed genes. Schematics illustrating the expression patterns of the target genes and the specific embryonic regions scanned for quantification. B and D) Visualizing transcriptional output. Representative MIP images generated from Z-stacks spanning the full depth of the cortical nuclear layer for the indicated genotypes. C and E) Quantitative impact of Fsh and dCBP depletion. Charts displaying the average fold change (FC) in smiFISH or HLB-RNAPII signals in fs(1)h or nej RNAi embryos relative to w RNAi controls. Fold changes were first normalized between embryos at matched developmental stages (cycles 12 and 13, determined by nuclear diameter) prior to calculating the mean FC for each gene.

The effects of dCBP (*nej*) knockdown mirrored those of Fsh knockdown but were markedly more severe (Fig. 9D–E). Depletion of dCBP nearly abolished the transcription of *sala, hb,* and *tld*. While we observed a slight reduction in *btd* transcription and RNAPII signal at the HLB, these effects were minor compared to the near-total loss of Zld-target expression. Collectively, these data demonstrate that while dCBP has a broader impact on the embryonic transcriptome, both dCBP and Fsh are essential for the robust transactivation of Zelda’s primary targets.

### Smr modulates the transcriptional output of Zld-target genes

We next investigated the corepressor Smr. RNA-seq in *smr* RNAi embryos revealed broad up-regulation of genes associated with the strongest Zld peaks (Fig. 10A, B). To specifically investigate the functional relationship between Smr and Zld, we associated Zld peaks with their nearest genes within a 15 kb window. We observed a positive correlation between Zld occupancy and Smr-mediated repression: genes associated with the strongest Zld peaks were significantly more likely to be up-regulated upon *smr* knockdown (Fig. 10B). This suggests that Smr plays a critical role in modulating Zld-dependent transactivation, potentially preventing premature or ectopic expression. This matches the presence of a conserved repression domain within the Zld protein [50].

**Fig. 10.**
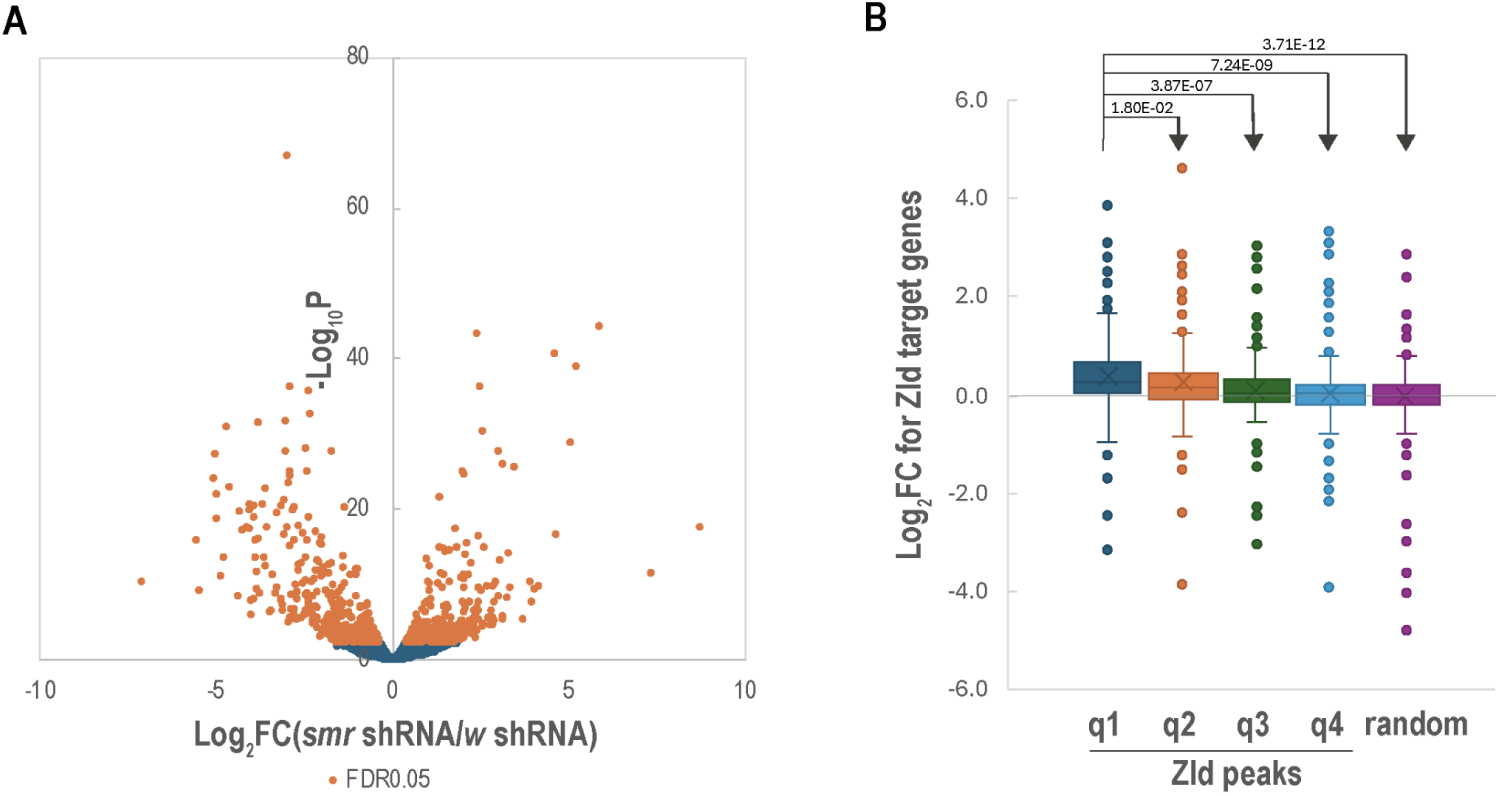
Depletion of Smr leads to the derepression of genes associated with strong Zld binding. A) Global transcriptomic changes upon smr knockdown. Volcano plot displaying differential gene expression in smr RNAi embryos (cycle 12–13) compared to w RNAi controls. Data were generated from RNA-seq and analyzed using the edgeR exactTest. Significantly up- and down-regulated genes are highlighted. B) Correlation between Zld occupancy and Smr-mediated repression. The top 3,000 Zld peaks were associated with their nearest neighbor genes. These peak-gene pairs were consolidated by assigning only the strongest Zld peak to each unique gene, resulting in a high-confidence dataset of 1,160 pairs. Genes were ranked by the intensity of their associated Zld peak and divided into four quantiles (q1–q4, with q1 representing the strongest Zld binding). Box plots show the distribution of fold changes in transcript levels following smr knockdown for each quantile, compared to a randomly selected set of control genes. P-values from a Student’s t-test are indicated above each quantile, demonstrating that genes with the strongest Zld occupancy exhibit the highest degree of derepression upon Smr depletion.

### Tlk interacts directly with Zld *in vitro*

Our IP-MS and CUT&RUN analyses indicated that the kinase Tlk is physically associated with Zld at genomic target sites. To determine if this interaction is direct, we performed *in vitro* GST pull-down assays. We partitioned Zld into four segments and expressed them as Glutathione S-Transferase (GST) fusion proteins. Immobilized fusion proteins were incubated with *in vitro* translated Tlk, and binding was assessed via Western blot.

As shown in Figure 11, untreated Tlk failed to bind any Zld segments. However, upon treatment with Calf Intestinal Phosphatase (CIP) to remove autophosphorylation or inhibitory phosphates, we observed significant and selective binding of Tlk to the Zld segment encompassing residues 401–750. A kinase-dead version of Tlk (Tlk^dead^) similarly exhibited robust binding to the same segment. These findings mirror the biochemical behavior of the nucleosome assembly factor Asf1, a known Tlk substrate [95,96], suggesting that Tlk’s kinase activity or its own phosphorylation state can autoinhibit its association with substrates. Collectively, these results suggest that Tlk directly interacts with Zld in a phosphorylation-dependent manner, providing a potential biochemical link between the cell-cycle machinery and the regulation of Zld activity during the rapid mitotic cycles of early ZGA.

**Fig. 11.**
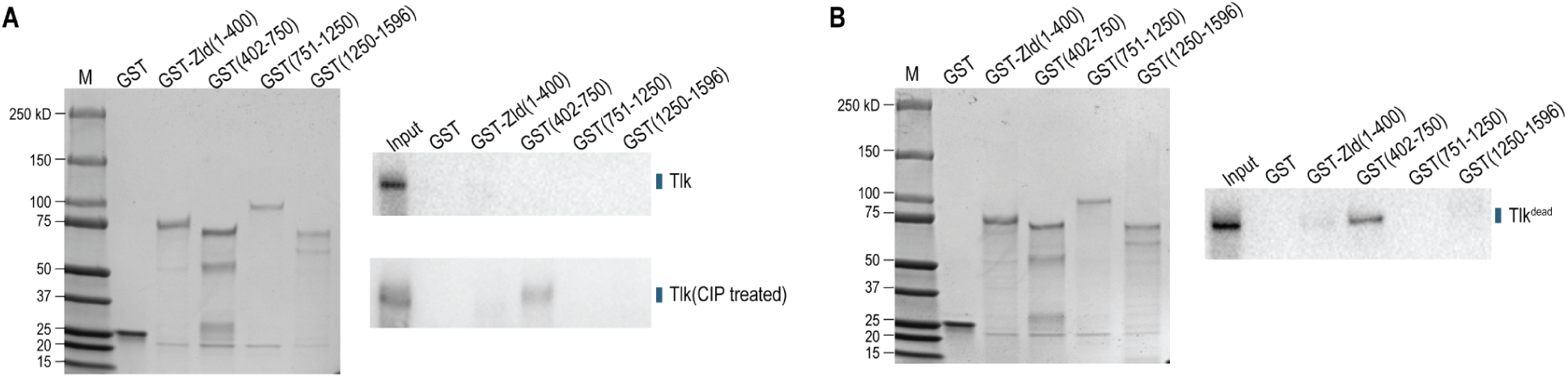
Direct interaction of Tlk with a Zld domain in vitro. GST-Zld fusion proteins containing indicated segments of Zld were immobilized on glutathione magnetic beads and incubated with in vitro translated Tlk protein, either wild-type (WT) or harboring a kinase-domain mutation (Tlkdead). Bound Tlk was analyzed by Western blot. **A)** Tlk specifically interacts with Zld(401–750). (Left) Coomassie-blue stained SDS-PAGE gel showing 20% of the immobilized GST or GST-Zld fusion proteins used as bait in the binding reactions. (Upper right) Western blot analysis of binding reactions with untreated WT Tlk. (Lower right) Western blot analysis of binding reactions with WT Tlk pre-treated with Calf Intestinal Phosphatase (CIP). The Input lane represents 20% of the total TNT product used per reaction. **B)** Mutation in the Tlk kinase domain enhances binding to Zld(401–750). Binding reactions performed as in (A), using the Tlk kinase-dead mutant as the prey protein. The Input lane represents 20% of the total TNT product used per reaction.

## DISCUSSION

In this study, we utilized an optimized IP-MS pipeline to define the diverse protein interactome of Zld. We identified multiple groups of functionally distinct factors, primarily those involved in chromatin remodeling and transcriptional control. Our functional validation of dCBP and Fsh extends recent reports of their roles in the MZT [85,87,97,98]. Furthermore, our identification of Smr provides a likely mechanistic explanation for Zld’s conserved repressive domain [50]. Finally, the discovery of a direct, phosphorylation-sensitive interaction between Zld and Tlk suggests a novel regulatory link between DNA replication and ZGA.

### Zld recruits specialized machinery to drive chromatin accessibility

Zld is uniquely required for nucleosome depletion and chromatin accessibility at zygotic enhancers [7,26–28,31,99], exerting a more profound effect on accessibility than other embryo patterning factors [26,31]. While the classical definition of a pioneer factor centers on the ability to bind motifs embedded in “closed” chromatin, recent evidence suggests this distinction is not absolute. Non-pioneer TFs can occasionally access closed chromatin depending on motif positioning [10,11,100,101], nucleosomal structures [102,103], and cellular concentrations [99,104,105]. Therefore, the hallmark of a pioneer factor should be defined not just by its occupancy of closed chromatin, but by its functional capacity to actively remodel the chromatin landscape.

Our IP-MS and CUT&RUN data provide a biochemical basis for this distinction. We demonstrate that Zld recruits a specific suite of SWI/SNF-family nucleosome remodelers - including PBAP/BAP and NURF—as well as the FACT complex to induce accessibility at its target sites. This partnership with chromatin remodelers is a conserved feature among high-potency pioneer factors across species [20,106]. In *Drosophila*, the pioneer factor GAF provides a primary example of this coordination, it has been shown to physically interact with NURF [107–109], PBAP [110], and ACF [43], as well as the FACT complex [111], to drive nucleosome remodeling and chromatin opening [68,112]. Similarly, mammalian pioneer factors such as Oct4, PU.1, and GATA3 physically recruit SWI/SNF (BAF) complexes to initiate chromatin opening, which in turn has been shown to stabilize the binding of the pioneer factors in some cases to sustain a permissive state [113–125].

Crucially, TFs appear to differ significantly in their inherent ability to recruit these remodeling machines [106,124]. While many early patterning factors may occupy the same enhancers, our results suggest that Zld acts as the primary “recruitment hub” for the remodeling apparatus. By integrating chromatin remodeling with the establishment of accessibility, Zld creates a permissive environment for other TFs that lack the specialized capacity to interface with the remodeling machinery. Our data reinforce a model where the “pioneer” status of Zld is defined by its role as a scaffold that bridges the gap between the closed chromatin state and the active transcriptional machinery.

### Recruitment of RNAPII by Zld to enhancers

A striking finding of our study was the robust RNAPII signal at distal Zld-bound enhancers. This contrasts with previous ChIP-chip [83] and ChIP-seq [84] studies where RNAPII was localized almost exclusively to promoters and gene bodies. We propose that RNAPII at enhancers may not be traditionally engaged with DNA in a pre-initiation complex, leading to the low cross-linking efficiency that rendered it “invisible” in prior ChIP assays.

Our results suggest that Zld co-occupies regulatory sites with the transcription machinery. In our imaging analysis, we observed spatial overlap between Zld and high-intensity RNAPII clusters, consistent with their functional dependence on Zld [88]. However, the overall degree of overlap was modest. This can be attributed to several biological and technical factors. First, the association is likely highly dynamic and occurs against a high stochastic background. Second, at cycle 12, less than half of RNAPII peaks at promoters are Zld-dependent [76]. Indeed, when we specifically analyzed the spatial association of Zld with the transcription foci of Zld-dependent genes (as detected by smFISH), we observed significantly higher overlap relative to rotated controls. Furthermore, transcription foci associated with large genes can exhibit low spatial colocalization with activators bound at distal enhancers or promoters [94,126].

The high density and uniform appearance of Zld clusters also present challenges for colocalization analysis. In contrast to Zld, patterning factors such as Dl [86] or Bicoid(Bcd) [35,36] tend to form significantly larger clusters. One likely explanation for this morphological difference is that Zld-target enhancers usually contain a relatively small number of binding sites, whereas Dl- and Bcd-target enhancers often possess dense clusters of their respective sites. Taken together, these biological and technical factors help to explain the modest overlap of RNAPII and Zld in our immunostaining analysis and likely account for the failure to detect this association in previous studies [35,37,87]. Additionally, we have demonstrated that the GFP-Zld lines used in these earlier reports contain fragmented proteins. This fragmentation would have increased diffuse background noise, further masking the subtle and dynamic colocalization signals that characterize these interactions.

The association of RNAPII with enhancers is a documented phenomenon, observed at ecdysone receptor-bound enhancers in *Drosophila* [127] and at enhancers and super-enhancers in other systems [128–131]. Enhancer-recruited RNAPII may be part of large transcription complexes [132] involved in eRNA transcription [128–130]. However, the failure of ChIP-based methods to detect these signals—despite robust detection by CUT&RUN—suggests that RNAPII at enhancers is primarily maintained through protein-protein interactions rather than the direct DNA-binding characteristic of pre-initiation or elongating complexes in the embryos. Consequently, we propose that RNAPII is recruited to Zld-bound enhancers to serve as a reservoir for the rapid assembly of transcription complexes at promoters, a mechanism recently proposed in other contexts [133]. This model echoes the behavior of pioneer factors Foxa3 and PHA-4/FoxA, which associate with paused RNAPII to facilitate rapid target gene expression [134,135]. This specialized function of Zld may be of particular importance for achieving the high-level transcriptional activation required within the extremely compressed mitotic windows of early development.

### dCBP and Fsh are critical for Zld-mediated transcriptional activation

Our functional analysis confirms that Zld initiates a regulatory cascade—Zld → dCBP → histone acetylation → Fsh—consistent with recent findings [85,87,97,98]. Fsh/BRD4 orthologs are known to bind acetylated histones to enhance transcriptional activation [52–54,81,82,91]. In our CUT&RUN experiments, we observed a strong correlation between H3K18ac and Fsh signals; similarly, imaging analysis revealed significant spatial overlap and a quantitative relationship dependent on the intensity of H3K18ac and Fsh clusters. Given this robust correlation and the fact that BRD4 binds H3K18ac (or H3K18ac/K23ac) with high affinity [82], H3K18ac likely plays a major role in Fsh recruitment, particularly given the strength of the H3K18ac signal around Zld peaks. While H3K27ac signals also correlate with Fsh in CUT&RUN, previous studies suggest it is not recognized efficiently by BRD4 [82], and is dispensable for Fsh recruitment in blastoderm embryos [136].

Interestingly, our CUT&RUN data reveal Fsh peaks directly overlapping Zld binding sites even where histone acetylation is depleted due to nucleosome displacement. This suggests that Fsh recruitment may also involve direct protein-protein interactions with Zld, potentially explaining why bromodomains or acetylation marks appear partially dispensable for activation in certain contexts [137]. In this regard, our anti-Zld IP-MS data suggest that the long isoform (Fsh-L) is more efficiently recruited to Zld binding sites, despite both isoforms sharing the bromodomains required for binding acetylated histones.

High-resolution imaging revealed hundreds of discrete, weak H3K18ac clusters as early as cycle 8, consistent with the low-level ChIP-seq signals detected at this developmental stage [27]. Both the number and intensity of these clusters increase consecutively through subsequent mitotic cycles. Because H3K18ac is maintained during mitosis (data not shown), this accumulation likely results from continuous de novo deposition. Consequently, our results suggest that Zld-driven ZGA is an incremental process characterized by the gradual accumulation of histone acetylation and a progressive increase in enhancer accessibility [30], which collectively culminate in the full activation of target genes.

### Interaction of Zld with chromatin boundary factors

Our identification of multiple chromatin boundary factors in the Zld interactome provides a biochemical basis for recent genomic studies. Zld motifs are highly enriched at Topologically Associating Domain (TAD) boundaries, and Zld is essential for the *de novo* formation of these boundaries during the MZT [71,72]. Furthermore, the BEAF-32 motif was previously identified as a top-ranking enriched sequence within Zld ChIP-seq peaks in both early embryos and larval neuroblasts [73].

Among these factors, Row is particularly noteworthy. While not a classical boundary protein, Row was enriched in both our anti-Zld and anti-GFP IP-MS experiments. Recent work has shown that Row interacts with BEAF-32 and other boundary factors to facilitate long-range interactions between regulated genes and TAD boundaries [138]. Interestingly, Row is also critical for neuronal development [139], a role that mirrors Zld’s function in maintaining neuroblast identity [73]. Together, our data strengthen the model that Zld organizes higher-order chromatin structure, a function that likely extends beyond early ZGA to influence cell-type-specific development in the nervous system.

### Coordination of Zld Activity with the cell cycle

Most intriguingly, the kinase Tlk—the most significantly enriched factor in our IP-MS—interacts directly with Zld via residues 401–750. Tlk is a highly conserved protein kinase whose activity is tightly coupled to the cell cycle, peaking during S-phase [70]. Notably, DNA replication has been shown to be required for Zld-mediated activation in early embryos [85]. Taken together, our observation that Tlk directly interacts with Zld suggests that Tlk provides a vital biochemical bridge, ensuring that Zld-driven ZGA is precisely coordinated with the rapid nuclear divisions in early embryos.

### Interaction of multiple coordinated factors with Zld

While our study identifies several critical factors involved in Zld activity, a recent study identified an additional cofactor, Ldb1, which acts as a bridge between Zld and dCBP [136]. We did not detect Ldb1 in our MS analysis, which may be attributed to its low abundance in our embryonic nuclear extracts or the inherent challenges in detecting certain proteins via mass spectrometry. Interestingly, neither Ldb1 [136] nor dCBP [98] appear to be required for Zld-induced chromatin accessibility, suggesting that Zld employs distinct sets of cofactors to manage different aspects of its pioneer function.

In summary, we have provided a comprehensive proteomic and genomic map of the Zelda interactome during the onset of ZGA. Our data reveal that Zld serves as a central hub, integrating the recruitment of nucleosome remodelers to open chromatin, the docking of RNAPII at distal enhancers to prime the transcriptional machinery, and the association of Smr to temper transcriptional output. Furthermore, the direct, cell-cycle-sensitive interaction between Zld and the Tlk kinase provides a potential biochemical link to coordinate these events with the rapid divisions of the early embryo. Collectively, these results elucidate the multi-layered mechanisms by which a master pioneer factor orchestrates the first major transition in embryonic development

## Materials and Methods

### Antibodies

The antibodies for Zld [140], Tlk [141], dCBP [142], Fsh [80], Smr [143], Brm and PB [110], Ssrp1 [111], NELF-A [144], and Med14 [145] have been described in previous studies, while the other antibodies including those against GFP (#1: Millipore, #AB3080P; #2: Thermo Fisher, #A11122), Fib (Abcam, #ab5821), H3K18ac (PTM Biolabs, #PTM-158), H3K27ac (Thermo, Catalog #MA5-23516), and RNAPII(clone: 4H8; Abcam, #ab5408) were purchased from commercial sources.

### Fly stock

Fly stocks
| Genotype | source | catalog no. |
| --- | --- | --- |
| w[*]; P{w[+mC]=matalpha4-GAL-VP16}V37 | BDSC | 7063 |
| y[1] v[1]; P{y[+t7.7] v[+t1.8]=TRiP.HMS00017}attP2 | BDSC | 33623 |
| y[1] sc[*] v[1] sev[21]; P{y[+t7.7] v[+t1.8]=TRiP.HMS01570}attP2/TM3, Sb[1] | BDSC | 36682 |
| y[1] sc[*] v[1] sev[21]; P{y[+t7.7] v[+t1.8]=TRiP.HMS02340}attP40 | BDSC | 41943 |
| y[1] sc[*] v[1] sev[21]; P{y[+t7.7] v[+t1.8]=TRiP.HMS00842}attP2/TM3, Sb[1] | BDSC | 34087 |
| zld294, FRT19A/FM7 | Liang et al., 2008 [23] |  |
| ovoD1,hsFLP112,FRT19A/Y | Liang et al., 2008 [23] |  |
| sfGFP-Zld | Hamm et al. 2017 [50] |  |

### IP - MS analysis for identification of Zld associated factors

#### Embryo collection

Wild-type (*Oregon-R*) or a line harboring endogenously GFP-tagged Zld [37] were used for IP-MS experiments. Flies were maintained in large population cages at 25°C. Embryos were collected for 1 h and allowed to develop for an additional 1 h 50 min at 25°C(to reach approximately cycle 12–14). Following harvest, embryos were dechorionated in 50% bleach for 3 min and rinsed thoroughly with water.

Embryos were resuspended in a formaldehyde-containing crosslinking buffer (15 mM HEPES, pH 7.6, 60 mM KCl, 15 mM NaCl, 4 mM MgCl_2_, 3% formaldehyde (Sigma-Aldrich, #F8775), 0.5mM DTT, and cOmplete protease inhibitor cocktail (Sigma-Aldrich, #4693159001)) and homogenized in a Dounce homogenizer (Pestle A, 7 strokes). The homogenate was incubated for 8 min at room temperature. Nuclei were pelleted by centrifugation at 2,500 rpm for 5 min (Eppendorf 5810R, S-4-104 rotor), resuspended in 10 mL PBST ( 137 mM NaCl, 2.7 mM KCl, 8 mM Na_2_HPO_4_, and 2 mM KH_2_PO_4_, 0.5% Triton X-100), supplemented with 0.125 M glycine, and 0.2 mM PMSF, and Dounce-homogenized again. After 5 min at room temperature, nuclei were pelleted at 3,000 rpm for 5 min, washed three times in PBST + 0.2 mM PMSF, and stored at -80°C.

#### Preparation of lysate from crosslinked nuclei

Nuclear lysates were prepared as previously described [41] with modifications. Crosslinked nuclei from 5 g of embryos were resuspended in 40 mL LB1 (50mM HEPES-KOH, pH 7.5, 140mM NaCl, 1mM EDTA, 20% glycerol, 0.5% NP-40, 0.25% Triton X-100, 1mM DTT, and 0.2mM PMSF), and homogenized in a dounce homogenizer (Pestle B, 20 strokes). After centrifugation at 2,500 rpm for 5 min, the pellet was washed in 20 mL LB2 (10 mM Tris-HCl, pH 8.0, 200 mM NaCl, 200 mM NaCl, 1 mM EDTA, 0.5 mM EGTA, and 0.2mM PMSF) and finally resuspended in 3.5 mL LB3(10 mM Tris-HCl, pH 8.0, 100 mM NaCl, 1 mM EDTA, 0.5 mM EGTA, 0.1% Na-deoxycholate, 0.5% N-lauroylsarcosine, and 0.2mM PMSF).

Samples were sonicated in 400 µL aliquots for 10 cycles (15 s ON/45 s OFF) using a Bioruptor (Diagenode) at medium power. Following sonication, Triton X-100 (1% final), MgCl_2_ (3 mM final), and Benzonase (100 U/mL) were added, followed by incubation at 30°C for 20 min. After centrifugation (10,000 × g, 10 min), the supernatant was collected. The pellet was re-extracted with 0.8 mL LB3, sonicated for 5 additional cycles, and the supernatants were combined. 5× RIPA buffer was added to a final 1× concentration, and the lysate was cleared by centrifugation at 20,000 × *g* for 10 min.

#### Immunoprecipitation (IP)reactions

For each IP, 0.35 ml of the nuclear lysate was diluted to 1 ml with RIPA buffer (50 mM TrisCl, pH 8.0, 0.15M NaCl, 0.1%SDS, 1% Triton X-100, 0.1% sodium deoxycholate, 1mM EDTA, 0.2mM PMSF), and incubated with 50 µL Protein A Dynabeads (ThermoFisher, #10002D) pre-bound with 8 µg of antibody: anti-Zld, anti-Fib, anti-GFP #1(Millipore, #AB3080P), or anti-GFP #2(ThermoFisher, #A6455). Reactions were incubated overnight at 4°C. Beads were washed consecutively: twice with RIPA, three times with RIPA + 1.0 M NaCl (or 2.0 M urea for high-stringency anti-Zld and anti-GFP #1 samples), three times with LiCl buffer (10 mM Tris-Cl, pH 8.0, 0.5 M LiCl, 1% NP-40, 1% Na-deoxycholate, 0.2 mM PMSF), and once with TE buffer.

#### Preparation of samples for mass spectrometry

Beads were eluted in 60 µL of the elution buffer (10 mM Tris-Cl, 1 mM EDTA, 200 mM NaCl, 1% SDS) at 99°C for 20 min, followed by a second elution in 30 µL. Proteins were reduced with 5 mM DTT (45°C, 30 min) and alkylated with 10 mM iodoacetamide (room temperature, 30 min, dark). Proteins were purified using the SP3 procedure [146] on paramagnetic beads and digested with 0.5 µg trypsin (Thermo Fisher, #90057)in 50 mM ammonium bicarbonate for 16 h at 37°C. Peptides were recovered, and concentrations were determined using Thermo Scientific Pierce Quantitative Colorimetric Peptide Assay Kit (#23275).

#### Mass Spectrometry analysis and data processing

Peptides were analyzed on a ThermoFisher Orbitrap Fusion Lumos Tribrid system interfaced with an Easy nLC 1200. Separation was performed on a C18 reverse-phase column (25 cm) using an organic gradient (5–30% ACN) over 180 min at 300 nL/min. Spectra were continuously acquired in a data-dependent manner throughout the gradient, acquiring a full scan in the Orbitrap (at 120,000 resolution with an AGC target of 400,000 and a maximum injection time of 50 ms) followed by as many MS/MS scans as could be acquired on the most abundant ions in 3 s in the dual linear ion trap (rapid scan type with an intensity threshold of 5000, HCD collision energy of 32%, AGC target of 10,000, maximum injection time of 30 ms, and isolation width of 0.7 *m/z*). Singly and unassigned charge states were rejected. Dynamic exclusion was enabled with a repeat count of 2, an exclusion duration of 20 s, and an exclusion mass width of ±10 ppm.

Data were analyzed using MaxQuant for label-free quantification (LFQ) [60] against the *D. melanogaster* proteome (FlyBase r6.29). Modifications Oxidation (M), Acetyl (Protein N-term), and Carbamidomethyl (C) were included in protein quantification. Matching between runs was enabled (0.7 min match window, 20 min time window). LFQ min ratio count was set at 1. All other parameters were set as default.

Perseus [61] was used for statistical enrichment. In the analysis, the filters for proteins include to have a minimum 2 valid values in each group. Imputation was used to replace missing values with parameters set at (1.8, 0.3) for the anti-GFP IP-MS data, but not for the anti-Zld and anti-Fib IP-MS data. For T-tests, *S*0 = 2 and FDR = 0.05 were applied, with a final enrichment cutoff of FDR 0.1. GO term enrichment was performed via the Gene Ontology online tool ( http://geneontology.org/) using the Fisher’s Exact method against the input sample reference list.

### CUT&RUN

#### CUT&RUN experimental procedures

Embryos were fixed and sorted as described in [97]. For each sample we used 75 sorted early (cycle 10–12) or late(cycle 12–13) stage 4 embryos. CUT&RUN experiments were conducted using a kit from Cell Signaling Technology (#86652S) with the buffer recipes following the original CUT&RUN protocol [147].

Briefly, fixed embryos were homogenized in a wash buffer (supplemented with spermidine and protease inhibitor cocktail). Nuclei were pelleted at 1500 × *g* for 5 min at 4°C, washed once, and then incubated with activated Concanavalin A magnetic beads for 20 min at room temperature on a rotator. Bead-bound nuclei were resuspended in the antibody binding buffer (supplemented with spermidine, protease inhibitors, and digitonin) and incubated with primary antibodies overnight at 4°C.

Following incubation, beads were washed twice with digitonin buffer and resuspended in 50 µL pAG-MNase pre-mix. After a 1 h incubation at 4°C, beads were washed twice. The pAG-MNase cleavage reaction was performed in 50 µL digitonin buffer supplemented with 1 µL of 100 mM CaCl_2_ for 1 h at 4°C. Reactions were stopped by adding 50 µL of 2×Stop Buffer (containing digitonin, RNase A, and yeast spike-in DNA) and incubated at 37°C for 10 min. To release DNA fragments, 1.5 µL of 20% SDS and 2 µL of 20 mg/mL Proteinase K were added, followed by overnight incubation at 65°C.

Samples were centrifuged at maximum speed for 5 min, and the supernatant was recovered on a magnetic rack. To enrich for short fragments, Ampure beads were added at a ratio of 65 µL per 100 µL of sample (0.65×). After 3 min, the beads (containing large DNA fragments > 600 bp) were discarded, and the DNA in the supernatant was recovered and purified using the Qiagen PCR purification kit. Sequencing libraries were prepared using the NEBNext Ultra II DNA Library Prep Kit (NEB, #E7645S) and barcoded with NEBNext Multiplex Oligos Dual Index Primers. Libraries were sequenced on an Illumina NextSeq P1 platform.

#### CUT&RUN data analysis

The 150 bp paired-end (PE) reads were mapped to the *D. melanogaster* genome (BDGP6) and the *S. cerevisiae* genome (R64-1-1; for spike-in normalization) using Bowtie2 with the following parameters: “--end-to-end --no-mixed --no-discordant -I 10 -X 700 -p 8 -S sample-name.sam”. Wig files were generated and normalized by scaling to 10 million aligned reads, followed by secondary normalization using the ratio of spike-in reads relative to the Zld sample. For visualization, signals were smoothed using a 0.3 kb window.

Peaks were called using MACS2 [148] with the following options: -c Input-DNA-reads -f BEDPE -g dm --min-length 500 --max-gap 500 --keep-dup all --call-summits. Inclusion of input DNA reads as control suppressed the calling of peaks over repeated sequences, but not entirely. To further eliminate artifacts from repetitive sequences, we used an IgG CUT&RUN sample as a control; any peaks showing a five-fold or greater enrichment in the IgG sample were discarded.

### Fly crosses for maternal RNAi knockdown

To achieve maternal knockdown, virgin females carrying *UASp-shRNA* transgenes were crossed to *Mat-tub-Gal4* males to drive shRNA expression during the later stages of oogenesis. Progeny siblings were intercrossed, and F1 progeny were grown at 27°C starting from the larval stage, to increase the Gal4/UASp induction [149]. Embryos were collected for 2.5 h, aged for 40 min, and fixed according to the protocol described by [150].

### Zld mutant germline clone generation

Maternal *zld* mRNA depletion was achieved using the FLP-DFS (dominant female sterile) technique [151,152] as previously described [27]. Briefly, *zld*^294^, *FRT19A*/*FM7* [23] virgin females were crossed with *ovoD1,hsFLP112,FRT19A*/Y [23] males. The larvae developed from embryos laid by females from these crosses were heat-shocked twice at 37°C for 2 h each (at 24–48 h and 48–72 h post-hatching) to induce mitotic recombination. Embryos from the resulting progeny were collected and fixed as described for the RNAi experiments.

### Immunofluorescence, smiFISH, and 3D co-localization analysis

#### Immunostaining

Embryos were collected from wild-type Oregon-R, *zld−/−* germline clones, pLEP_*hb*-stripe, pLEP_*hb*-stripe(zm), or RNAi knockdown flies (see below) and fixed as previously described [150]. For each sample, 20 µL were rehydrated, washed in PBST buffer (1×PBS buffer + 0.3% Triton X-100), and blocked for 45 min at room temperature in Western

Blocking Reagent (Sigma Aldrich, #11921673001) diluted in 1×PBS. Primary antibody incubation was performed overnight on a rocker at 4°C in 200 µL blocking solution using the following antibodies at recommended dilutions: anti-Zld, anti-Fsh[80], anti-H3K18ac (PTM Biolabs, #PTM-158), anti-RNAPII (4H8) (Abcam, #ab5408), and anti-GFP (Aves Labs, Inc, #1010).

The embryos were briefly washed three times with PBST, followed by six washes of 20 min each (1 mL PBST per wash). After a second blocking step, embryos were incubated with Alexa Fluor-conjugated secondary antibodies (647, 555, or 488; Thermo Fisher), diluted as recommended in PBST for 2 hrs at room temperature. After the incubation, the embryos were washed five times for 20 min each with PBST. Nuclei were stained with Hoechst 33342 (Thermo Fisher, #H3570) at the concentration of 1 µg/ml for 30 min. Embryos were mounted in ProLong™ Diamond Antifade Mountant (Thermo Fisher, #P36961) (60 µL suspension in mountant per slide) and cured for 24 hrs in the dark before imaging.

#### smFISH

To analyze colocalization of Zld clusters with transcription foci of *zen, dfd, and lacZ*, immunostaining with anti-Zld antibody was performed following smiFISH as described in Calvo et al. [153]. For smiFISH, 20 nt gene-specific primary probes used for the assay (42 for *zen*-RA, 36 for *dfd*-RA, and 49 for *lacZ*) were designed using the Stellaris® RNA FISH Probe Designer (Biosearch Technologies, Inc.), and synthesized by IDT(Integrated DNA Technologies); all probe sequences are provided in Supplementary Table 3. Each probe included a 5′ common sequence (CCTCCTAAGTTTCGAGCTGGACTCAGTG) for annealing to a X-FLAP oligo (CACTGAGTCCAGCTCGAAACTTAGGAGG) labeled at both 5′ and 3′ ends with Alexa Fluor® 488 to allow detection.

For detection of the RNA of *sala*, *hb*, and *tld* in RNAi knockdown embryos, the smiFISH method was employed, with the use of a set of 20 probes for *sala-RA*, 43 for *hb-RA*, and 40 for *tld-RA*, respectively. X-FLAP oligo was labeled at both 5′ and 3′ ends with Alexa Fluor® 488(*hb*) or 647(*sala* and *tld*). For *btd*, smFISH using Custom Stellaris® FISH probes (Biosearch Technologies, Inc.) labeled with Quasar 670 was performed following the standard smFISH method (Biosearch Technologies, Inc.). For co-detection of RNAPII, smFISH was followed by standard immunostaining using the RNAPII 4H8 antibody (1:400) and Alexa Fluor 555-conjugated secondary antibody (1:1000). Samples were mounted in ProLong™ Diamond Antifade as described above.

#### Image acquisition

High-resolution imaging was performed on a Zeiss LSM800 Airyscan microscope using a Plan-Apochromat 63×1.4 NA oil DIC M27 objective. Images were acquired in Airyscan SR mode. The standard acquisition settings were as follows: pixel size of 0.035 µm, z-slice thickness of 0.13 µm, frame scan mode, 16-bit depth, line averaging of 2, digital gain of 2, master gain of approximately 850V. Laser power, digital gain, and master gain were adjusted such that the peak signal intensity was approximately 25% of the saturation level. Experiment-specific variations to these settings are detailed below.

For Zld and RNAPII colocalization analysis (Fig. 6), a 6× digital zoom and a scan speed of 5 were used. Laser power was set to 0.1% for both Alexa Fluor 555 and 488 channels. The image size was 16 µm × 16µm(454 × 454 pixels). and fields of view were selected from the central region of the embryos. To analyze the effects of *zld* mutation and *nej* RNAi knockdown on H3K18ac and Fsh cluster formation(Fig. 8), settings were identical except for a scan speed of 6, a digital gain of 1, and laser powers of 0.2% (Alexa 555) and 0.1% (Alexa 488).

For colocalization analysis of Zld with target gene transcription foci(Fig. 7), the zoom was 2× with a scan speed of 5. Laser powers were 1% for smFISH channels and 0.3% for the anti-Zld immunofluorescence channel. The image size was 50 µm × 50 µm(1412 × 1412 pixels). Images for *zen* and *dfd* were acquired from the dorsal-anterior region where their expression domains overlap. The *lacZ* imaging was performed over the posterior stripe controlled by the *hb* stripe enhancer.

To analyze the effects of *fs(1)h* and *nej* RNAi knockdown on transcription of Zld target and non-target genes, a 1.4× zoom and a scan speed of 6 were used for *btd* smFISH in *fs(1)h* RNAi and its corresponding w RNAi control embryos. Other images were acquired with a 1.0× zoom and scan speed of 5. Laser powers were 1% for all smFISH channels and 0.1% for RNAPII. The image size was 50 µm × 72 µm (1400 × 2028 pixels) for *btd*, and 25 µm × 100 µm(688 × 2850 pixels) for all others. Imaging for *hb*, *sala*, and RNAPII was centered around 30%–50% egg length(EL). For *btd*, imaging was centered at approximately 30% EL (a 75 µm × 25 µm section centered on the *btd* stripe was subsequently cropped for analysis). For *tld*, the imaging area was centered at 25 µm toward the dorsal side at the embryo midline.

#### 3D Object-based co-localization analysis

Image processing was performed in Fiji. To eliminate background, signals outside the nuclear boundaries in each Z-slice were removed using the “Clear Outside” function. Transcription foci (typically 1–2 per nucleus) (for *zen*, *dfd*, and *lacZ*) were identified as distinct 3D maxima using Fiji following background subtraction; intensity cutoffs were established for each gene to distinguish signal from noise and remained consistent across all images in an experiment. Nuclear spots from immunostaining images (anti-Zld, RNAPII 4H8, H3K18ac, Fsh, anti-GFP for GFP-Zld) were segmented using the Trainable Weka Segmentation 3D plugin. The resulting classified image stacks were converted into masks and applied to the original images using the 3D Object Counter. To identify local signal maxima within segmented objects, a 3D Maxima Filter plugin was applied to the masked images. For the Fsh-H3K18ac analysis, we further refined the segmentation by applying dilate and watershed functions to the classified masks to ensure the separation of fused clusters; this yielded fewer but more spatially distinct spots. For the Zld-RNAPII analysis, a one-slice Z-shift between the Alexa 555 and Alexa 488 channels was corrected prior to image processing.

Co-localization was quantified by calculating the Euclidean distance from each peak in the source channel to the nearest peak in the target channel. The search radius included the immediate z-slice and adjacent slices (assuming a z-dimension distance of 0.067 µm, half the slice thickness). To ensure we analyzed meaningful protein clusters rather than single molecules or background, we applied an intensity cutoff; only peaks with intensities 1.5× to 2× the average intensity of single-molecule clusters (identified as small, uniform-intensity spots at metaphase) were included.

To statistically validate colocalization, we utilized a 90-degree rotation control. The target channel image and its associated peak coordinates were rotated by 90 degrees, and Euclidean distances to the reference channel were recalculated. To ensure a fair comparison, only peaks that remained within the boundaries of the rotated image frame were included in the final analysis. This approach provides a more stringent control for stochastic overlap than random-coordinate assignment, as it preserves the original spatial architecture and cluster density while specifically disrupting biological coordination.

#### Image analysis for the RNAi experiments

Individual RNA molecules and transcription foci (typically 1–2 per nucleus) were identified as distinct 3D maxima using Fiji following background subtraction, Intensity cutoffs were established for each gene to distinguish signal from noise and remained consistent across all images in an experiment. RNAPII clusters corresponding to the Histone Locus Body (HLB) were analyzed by applying a Gaussian filter followed by segmentation with the Trainable Weka Segmentation 3D plugin. Masks generated from classified images were processed using the “Analyze Particles” and “Measure” functions to determine cluster intensities.

### RNA-seq in Smr RNAi knockdown embryos

Embryos from RNAi knockdown flies for the *smr* gene or the *white* (*w*) gene (control) were generated and collected as described above. Three biological replicates were prepared for both the *smr* RNAi and *w* RNAi groups, using 50 sorted Stage 4 embryos per sample. Total RNA was extracted using TRIzol (Thermo Fisher Scientific) and treated with TURBO DNase I.

Library preparation and sequencing were performed by the QB3-Berkeley Genomics core facility. Briefly, poly(A) mRNA was enriched using poly-dT beads, and libraries were constructed using the KAPA mRNA HyperPrep kit (Roche, #KK8581). Truncated universal stub adapters were ligated to cDNA fragments, followed by PCR extension using unique dual-indexing primers to generate full-length Illumina adapters. Libraries were pooled by molarity and sequenced on an Illumina NovaSeq X (10B flow cell) for 2 × 150 cycles.

Sequencing reads were mapped to the *D. melanogaster* genome (r6.59) using the STAR (v2.7.11b) aligner [154]. RNA-seq aligner to map the resulting reads to the *D. melanogaster* genome (r6.59). Gene-level coverage was generated from the resulting BAM files using featureCounts [155]. Differential expression analysis was performed using the exactTest method within the edgeR pipeline [156] to identify genes significantly affected by *smr* knockdown.

To investigate the specific regulatory link between Zld and Smr, Zld peaks from CUT&RUN experiments were assigned to the nearest gene within a 10 kb window. For genes associated with multiple Zld peaks, only the peak with the highest signal intensity was retained. Zld peaks were then partitioned into quantiles based on intensity. We analyzed the distribution of log_2_FC values for genes within each quantile to assess the relationship between Zld binding strength and Smr-dependent regulation. As a control, log_2_FC distributions were compared against randomly selected gene sets of equal size drawn from the complete edgeR output.

### In vitro protein binding assay for Tlk and Zld interaction

To generate protein expression plasmids, DNA sequences encoding specific Zld segments (aa 1–400, aa 401–750, aa 751–1250, and aa 1251–1596) were cloned into pGEX-6P-1-H-RBS vector (GenScript). The GST-fusion proteins were expressed in *E. coli* (NEB, #C3037) and purified using Pierce™ Glutathione Magnetic Agarose Beads (Thermo Scientific, catalog#: 78601) according to the manufacturer’s instructions. Briefly, bacterial lysates were prepared in lysis buffer (20 mM Tris-HC1 at pH 8.0, 0.2 M NaC1, 20 μM ZnC1_2_, 1 mM DTT, 0.5 mM phenylmethylsulfonyl fluoride (PMSF), and cOmplete protease inhibitor cocktail (Sigma-Aldrich, #4693159001)) and incubated with the magnetic beads. Following extensive washing, the GST-fusion proteins remained coupled to the beads for use in the binding assays. Protein concentrations on the beads were estimated by SDS-PAGE analysis of 5 µL bead aliquots alongside a bovine serum albumin (BSA) standard curve.

Tlk in vitro transcription/translation plasmid was constructed by inserting the Tlk encoding sequence into a pTnT Vector (Promega, cat no. L5610). Tlk protein was generated using the TnT® Coupled Reticulocyte Lysate System (Promega, #L4610) following the manufacturer’s protocol.

For the binding assay, approximately 3 μg of the purified GST or GST-fusion proteins immobilized on glutathione-agarose beads were incubated with 5 µL of the TnT product. The reaction was performed in 0.6 mL of protein-binding buffer (20 mM HEPES at pH 7.6, 100 mM KC1, 20 μM ZnC1_2_, 2.5 mM MgC1_2_, 10% glycerol, 0.025% NP-40, 0.5 mM DTT, 0.5 mM PMSF) supplemented with 600 µg of BSA to minimize non-specific interactions. After incubation at 4°C for 2 h with continuous mixing, the beads were washed four times with 1 mL of binding buffer. Bound Tlk protein was eluted in SDS sample buffer, resolved by SDS-PAGE, and detected by immunoblotting with an anti-Tlk antibody.

## Acknowledgements

We would like to thank members of the Eisen lab for helpful discussion and inputs throughout the project, and Drs. Susumu Hirose, Alexander Mazo, Jenn-Yah Yu, Ong Chin Tong, and Nicola Iovino for generous gifts of antibodies; We thank Lori Kohlstaedt of the Proteomics/Mass Spectrometry Laboratory at the University of California at Berkeley for mass spectrometry, and the QB3 Genomics, UC Berkeley, Berkeley, CA, RRID:SCR_022170 for RNA-seq library preparation and the next generation sequencing services. This was funded by an HHMI Investigator award to MBE.

## Author contributions

X.Y.Li and M.B.Eisen conceived the project. X.Y.Li performed all the experiments and data analysis. M.B. Eisen supervised the project. X.Y.Li and M.B.Eisen prepared the manuscript.

## Data availability

Raw imaging data and custom Python scripts for object-based colocalization analysis and for generating cluster peak-centered heatmaps are available at Zenodo (https://doi.org/10.5281/zenodo.19489735).

---

**Fig. 1—figure supplement 1.**
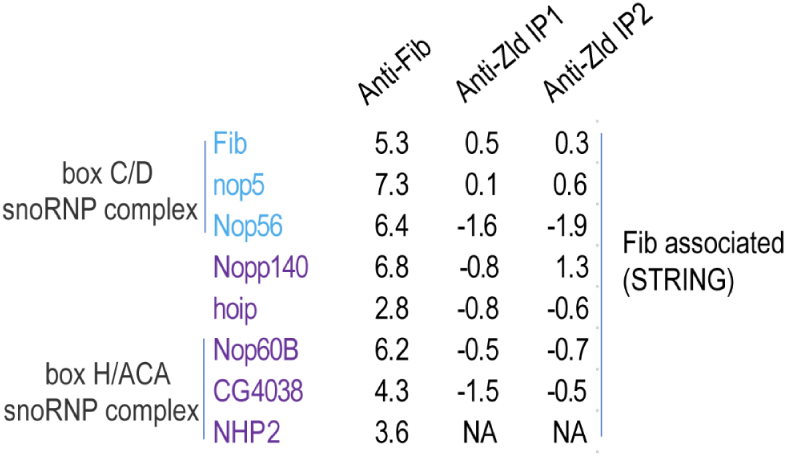
Enrichment of protein factors predicted to interact with Fib in anti-Fib IP-MS experiment. The enrichment values (log2(IP/Input)) for the seven factors that are among the top 10 predicted Fib associated factors in the STRING database [63] in the anti-Fib and anti-Zld IP-MS experiments are shown.

**Fig. 3—figure supplement 1.**
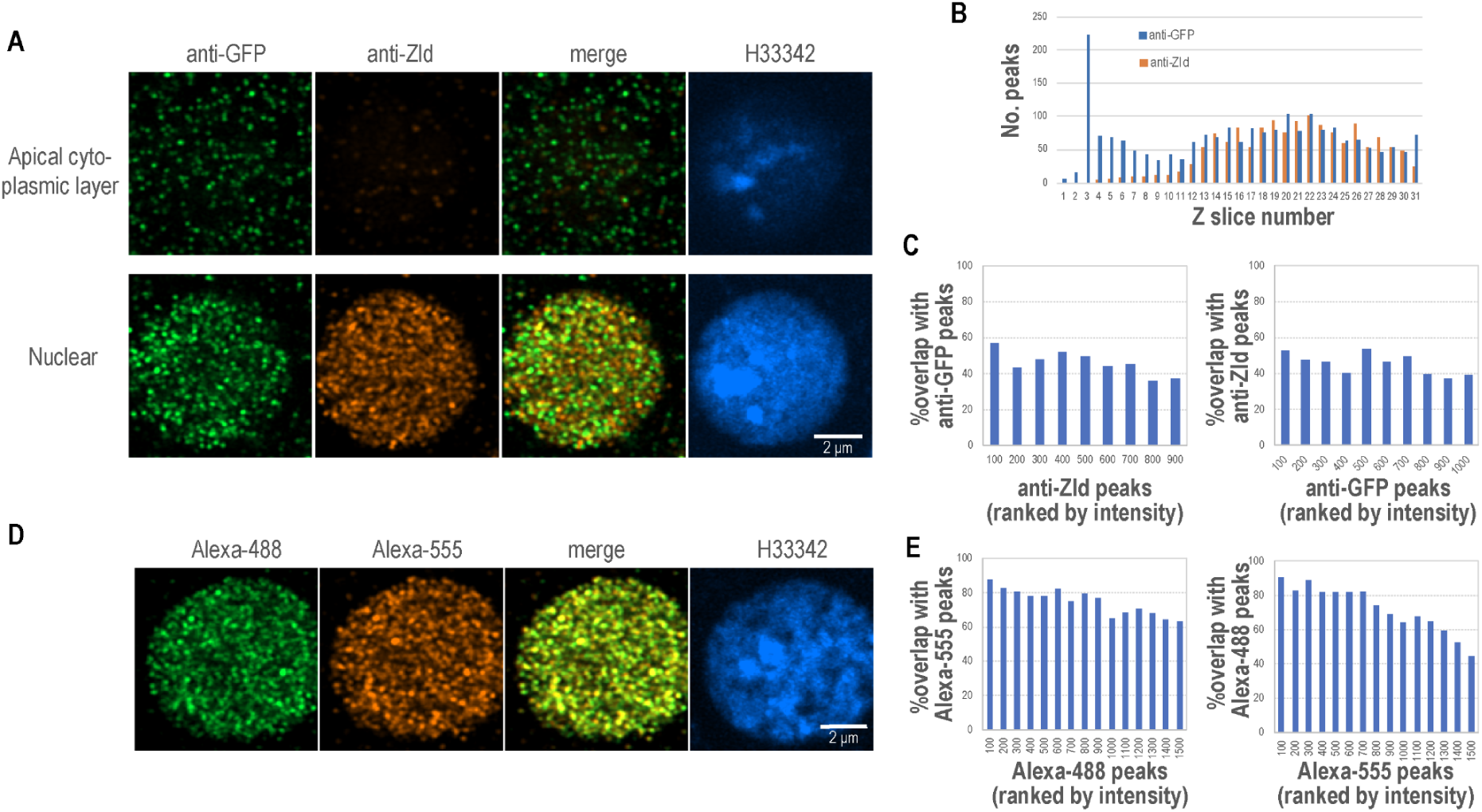
In vivo evidence of GFP-Zld fragmentation and spatial segregation. Embryos were stained with the indicated antibodies and high-resolution images were acquired using the Zeiss Airyscan detection system. Images were processed and analyzed using Fiji software as detailed in the Methods. A) Spatial segregation of Zld fragments in the cytoplasm and nucleus. Maximum intensity projections (MIP) of Z-slices (n = 2) from either the apical cytoplasmic layer or the nuclear center. In the cytoplasm, immunofluorescent signals were detected exclusively by the anti-GFP antibody. The absence of anti-Zld signal (targeting the C-terminal DNA binding domain) indicates these are isolated N-terminal fragments. In the nuclei, signals were detected by both antibodies. B) Z-profile of Zld fragments. Distribution of fluorescent signals detected by anti-GFP and anti-Zld antibodies across the full Z-stack, illustrating the relative localization of fragments throughout the embryo depth. C) Quantitative colocalization analysis of nuclear Zld fragments. Frequency of colocalization between foci detected by anti-GFP and anti-Zld antibodies. The distance from each spot in one channel to the nearest spot in the other was calculated; the chart shows the percentage of spots colocalizing within a 0.14 µm cutoff, binned by cohorts of 100 peaks ranked by intensity. The relatively low overlap demonstrates that the Zld fragments, not just full length Zld, are present in the embryo nuclei. D) Technical control for colocalization efficiency. Wild-type embryos were incubated with a single primary anti-Zld antibody followed by two secondary antibodies conjugated to different fluorophores. MIP images for both secondary antibody channels are shown. E) Validation of colocalization methodology. Quantification of the overlap between the two secondary antibody channels from (D), using the same proximity-based analysis described in (C). The high overlap serves as a baseline for maximum achievable colocalization.

**Fig. 3—figure supplement 2.**
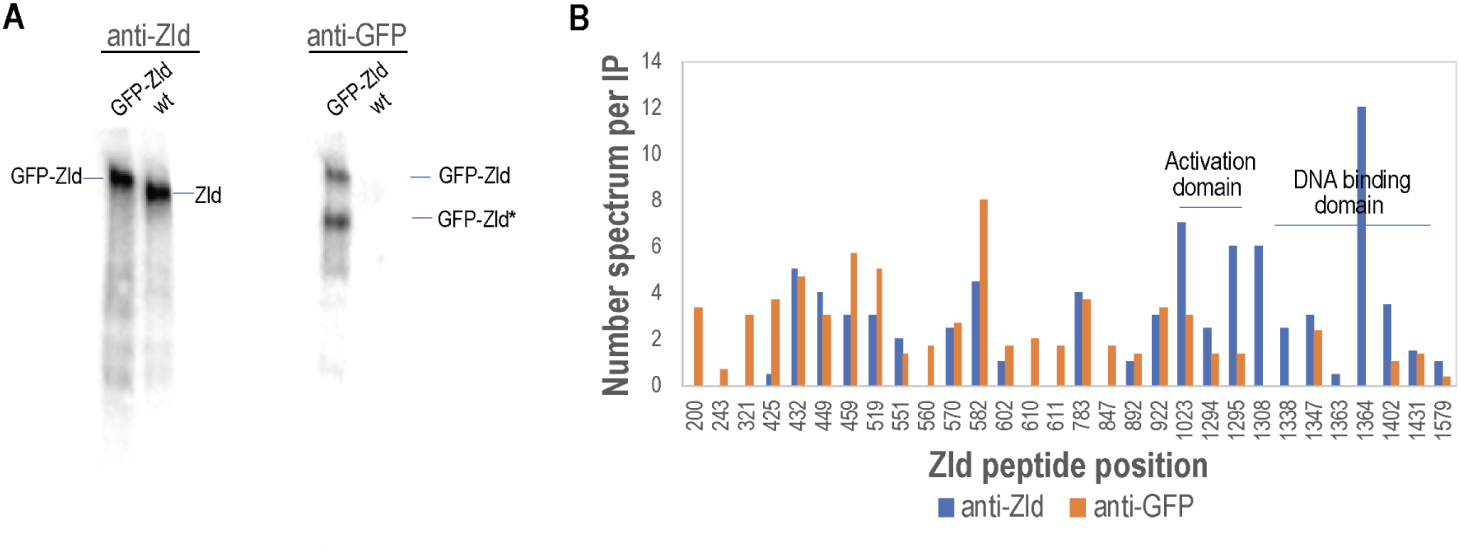
Evidence of GFP-Zld fragmentation in embryonic lysates. A) Detection of full-length and truncated GFP-Zld by Western blot. Full-length GFP-Zld is detected by the anti-Zld antibody (targeting the C-terminus), whereas both the full-length protein and a truncated N-terminal fragment (GFP-Zld*) are detected by the anti-GFP antibody. B) Peptide distribution across the Zld protein sequence. Distribution of peptide spectral counts across the Zld primary sequence as identified in anti-Zld and anti-GFP IP-MS experiments. The sharp decrease in spectral counts in the anti-GFP samples corresponds to the C-terminal region. The putative activation domain and the DNA binding domain (zinc-finger cluster) [49] are indicated for reference.

**Fig. 4—figure supplement 1.**
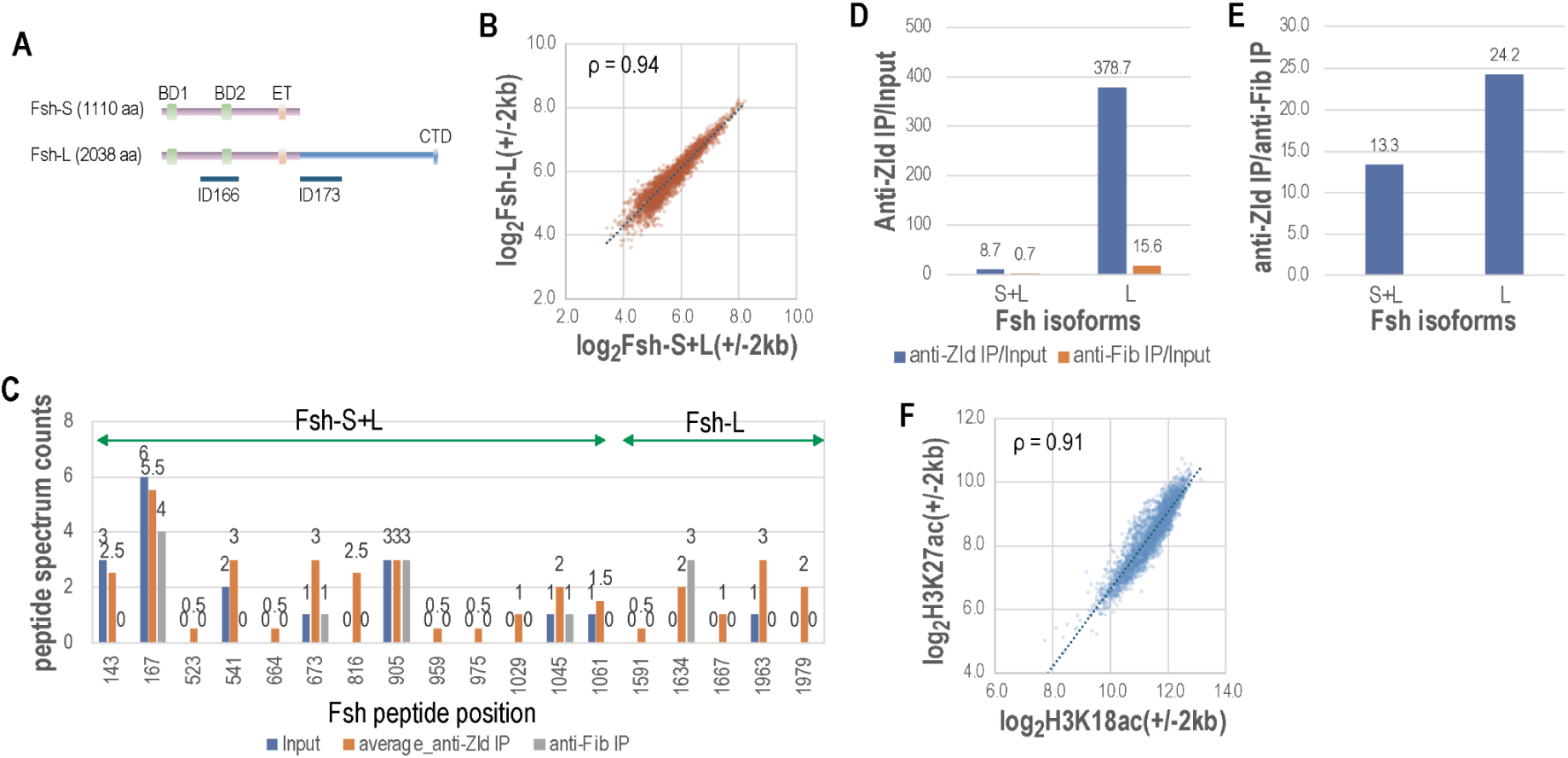
Enrichment of Fsh-L in anti-Zld IP-MS and CUT&RUN. **A)** Schematic structures of Fsh isoforms. Conserved domains and antigen regions for antibodies ID166 (recognizing both isoforms) and ID173 (isoform L-specific) are indicated (Kockmann et al. 2013). **B)** High correlation of CUT&RUN signals between the two Fsh antibodies at Zelda binding sites. ρ denotes the Spearman rank correlation coefficient. **C)** Fsh peptide positions and spectrum counts from anti-Zld IP-MS. Horizontal bars indicate peptides shared by both isoforms (Fsh-S+L) versus those unique to the long isoform (Fsh-L). D–E) Relative enrichment of Fsh-S+L and Fsh-L in anti-Zld IPs compared to Input (D) or anti-Fib control IP (E). Enrichment was calculated as the ratio of total peptide intensities in anti-Zld IPs to the corresponding intensities in the control samples. F) High correlation between H3K18ac and H3K27ac CUT&RUN signals across the genome. ρ denotes the Spearman rank correlation coefficient.

**Fig. 6—figure supplement 1.**
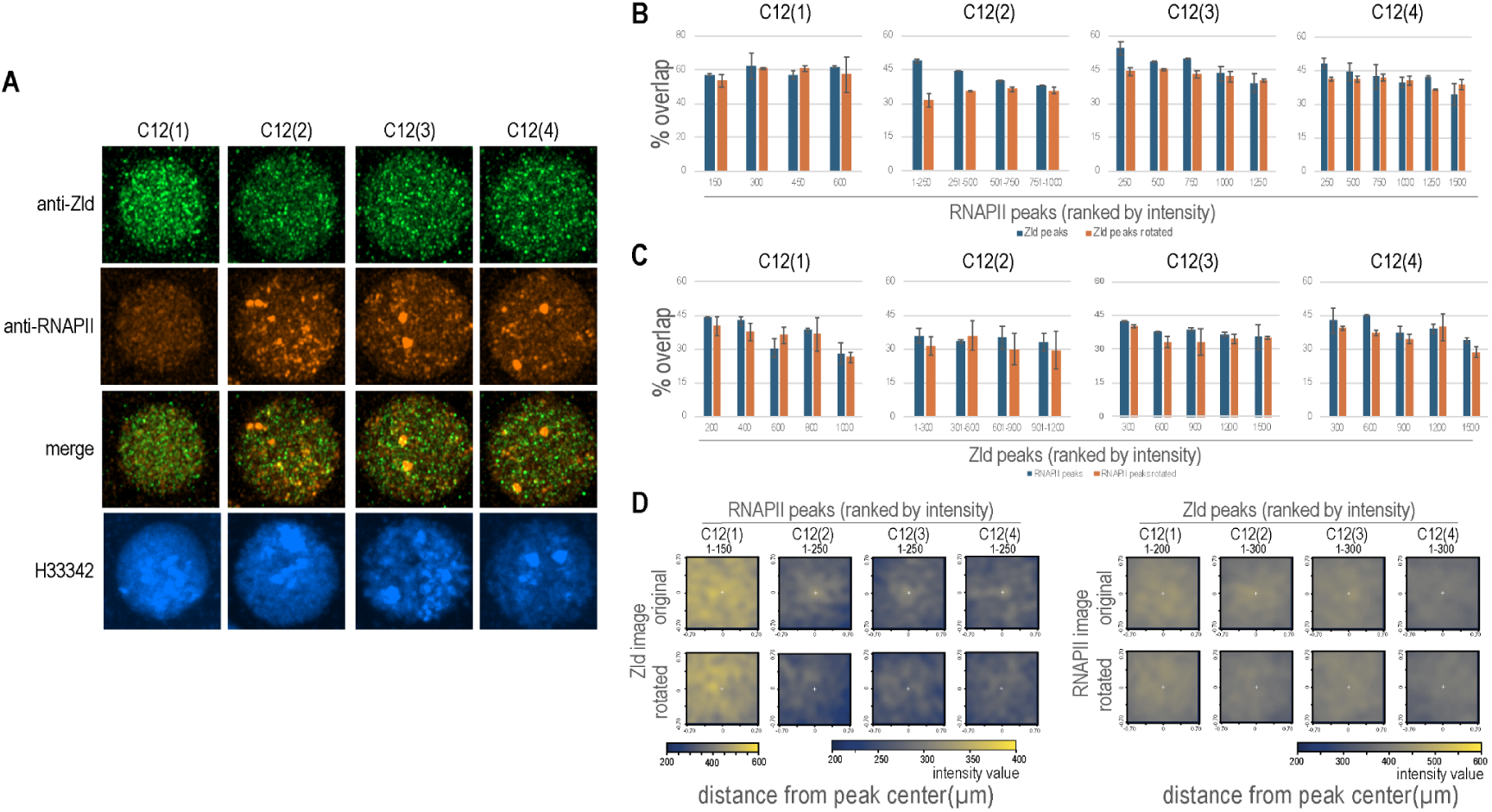
Temporal dynamics of Zld–RNAPII colocalization during cycle 12. High-resolution 3D colocalization analysis was performed across sequential stages of nuclear cycle 12 (C12.1 to C12.4). A) Temporal progression of transcriptional clusters. MIP images of the whole nuclei from embryos at different stages in mitotic cycle 12. B) RNAPII-centric colocalization. Percentage of RNAPII peak overlap with Zld (0.14 µm cutoff). Preferential association of the brightest RNAPII hubs is highest at C12.2 and diminishes at later stages as cluster intensities weaken; no significant enrichment is observed at C12.1 when RNAPII clusters are nascent. C) Zld-centric colocalization. Percentage of Zld peak overlap with RNAPII (0.14 µm cutoff). In contrast to RNAPII-centric analysis, enrichment over rotated controls is minimal across all cohorts and stages. D) Signal intensity heatmaps. Average Zld signal centered on top-ranked RNAPII cohorts and RNAPII signal centered on top-ranked Zld cohorts. Consistent with object-based analysis, Zld enrichment at RNAPII hubs is visible from C12.2 to C12.4, whereas RNAPII enrichment at Zld peaks is negligible, with only modest signal observed at C12.2.

**Fig. 8—figure supplement 1.**
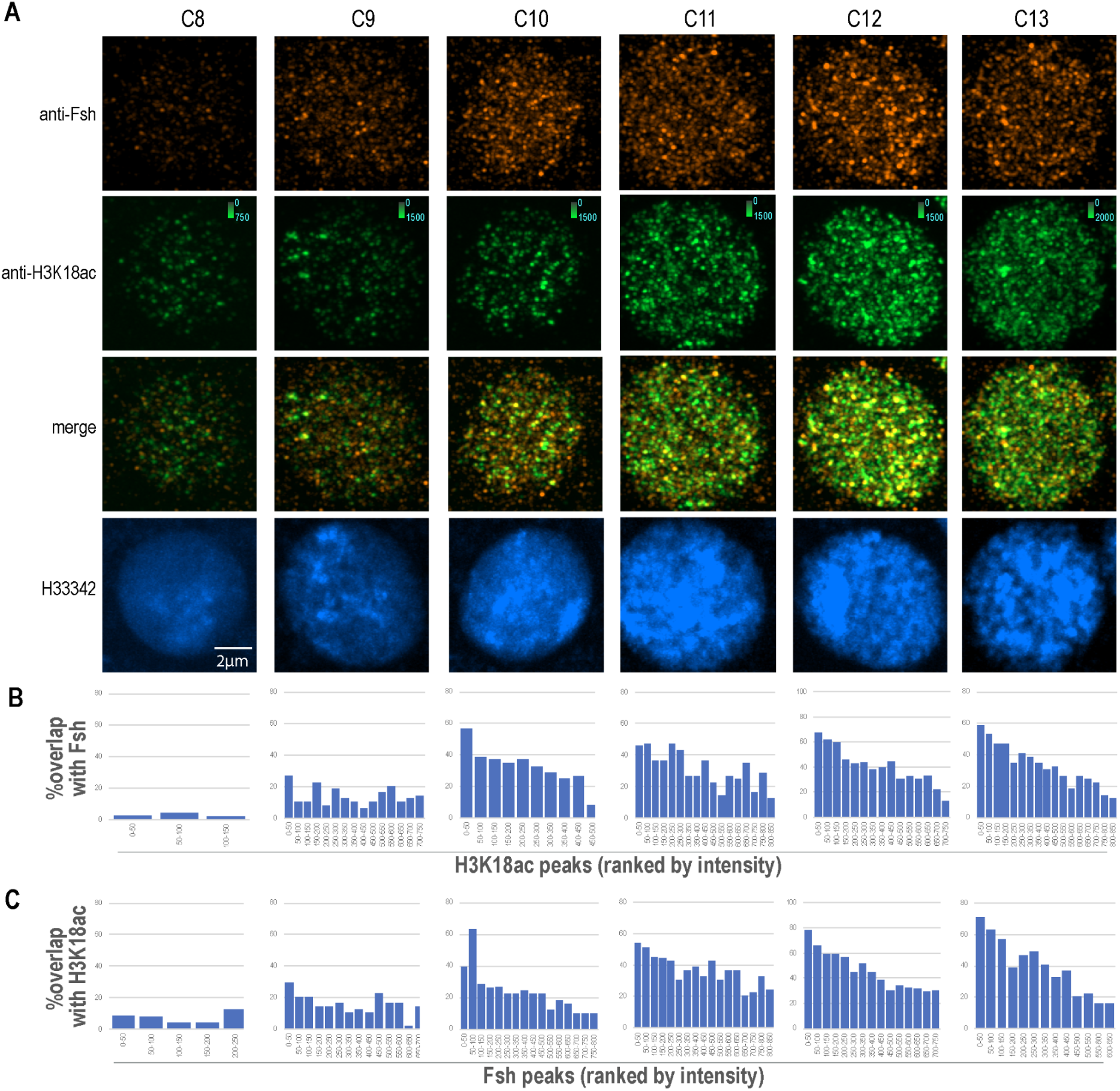
Dynamics of H3K18ac and Fsh clusters in early mitotic cycles. Similar to the main figure Fig.8. A) Developmental trajectory of H3K18ac and Fsh clusters. Representative MIP images of individual nuclei from embryos at the indicated mitotic cycles (C8–C14). Embryos were co-immunostained with anti-H3K18ac and anti-Fsh antibodies, illustrating the progressive increase in cluster number and intensity over developmental time. B) Intensity-dependent colocalization of Fsh with H3K18ac. Quantitative analysis showing the percentage of Fsh peaks that colocalize with H3K18ac peaks (within a 0.14 µm center-to-center cutoff). Fsh peaks were ranked by intensity and divided into cohorts of 50. The data demonstrate that stronger Fsh clusters are more likely to be associated with H3K18ac-enriched regions. C) Intensity-dependent colocalization of H3K18ac with Fsh. Quantitative analysis showing the percentage of H3K18ac peaks that colocalize with Fsh clusters (within a 0.14 µm cutoff). H3K18ac peaks were ranked by intensity and divided into cohorts of 50. Similar to (B), higher levels of histone acetylation correlate with a higher frequency of Fsh recruitment.

**Fig. 8—figure supplement 2.**
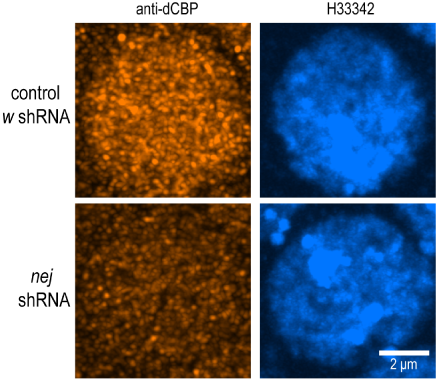
RNAi knockdown of dCBP. Immunostaining was performed with an anti-CBP antibody in embryos that expressed shRNA for the nej gene or the w gene (negative control). MIP images of individual nuclei are shown.

**Fig. 9—figure supplement 1.**
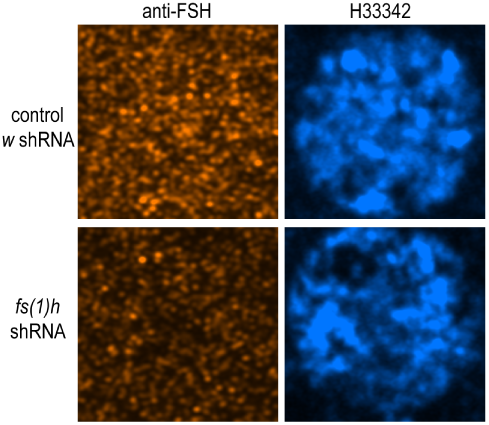
RNAi knockdown of Fsh. MIP images from immunostaining with anti-Fsh antibody in an embryo that expressed shRNA for the fs(1)h gene or the w gene (control)

